# Beta-cell Metabolic Activity Rather than Gap Junction Structure Dictates Subpopulations in the Islet Functional Network

**DOI:** 10.1101/2022.02.06.479331

**Authors:** Jennifer K. Briggs, Vira Kravets, JaeAnn M. Dwulet, David J. Albers, Richard K. P. Benninger

## Abstract

Diabetes is caused by dysfunction of electrically coupled heterogeneous *β*-cells within the pancreatic islet. Functional networks have been used to represent cellular synchronization and study *β*-cells subpopulations, which play an important role in driving dynamics. The mechanism by which highly synchronized *β*-cell subpopulations drive islet function is not clear. We used experimental and computational techniques to investigate the relationship between functional networks, structural (gap-junction) networks, and underlying *β*-cell dynamics. Highly synchronized subpopulations in the functional network were differentiated by metabolic dynamics rather than structural coupling. Consistent with this, metabolic similarities were more predictive of edges in the islet functional network. Finally, removal of gap junctions, as occurs in diabetes, caused decreases in the efficiency and clustering of the functional network. These results indicate that metabolism rather than structure drives connections in the function network, deepening our interpretation of functional networks and the formation of functional sub-populations in dynamic tissues such as the islet.

## Introduction

Diabetes Mellitus is a global epidemic afflicting > 500M adults world-wide and is associated with dysfunction or death of insulin-producing *β*-cells within the islets of Langerhans, located in the pancreas. *β*-cells are functionally heterogeneous and electrically coupled through connexin36 (C×36) gap junctions^1–3^, which allows insulin to be released in coordinated pulses^1,4–6^. Both the heterogeneity and electrical coupling are central to regulating the homeostasis of glucose and other nutrients^4,7–9^. In diabetes, there can be both an insufficient number of adequately functioning *β*-cells^7–10^ and lack of intra-islet *β*-cell communication^8,11,12^. Therefore, the role of *β*-cell heterogeneity in islet communication and collective synchronization is fundamental to understanding diabetes.

Network analysis is a promising tool for investigating the role of functional heterogeneity in biological systems. Network analysis can quantify the relationships between interacting parts of a system^13–17^; represented by its entities (nodes) and their interactions (edges). In the islet, nodes can correspond to *β*-cells, while edges represent either *structural* or *functional* relationships^17–20^. *Structural networks* represent physical connections, such as gap junctions. Obtaining a complete structural network of a biological system requires 3D imaging with techniques that differentiate cells and their connections. In dynamic tissues such as the islet, the structural network alone cannot describe functional behavior, which emerges from complex interactions between individual cell dynamics and the underlying structure^14,21^. *Functional networks* represent the emergent system behavior, where edges correspond to functionally similar cell pairs, for example defined by correlated [Ca^2+^] dynamics^17,21–23^. In diabetes the islet shows disrupted [Ca^2+^] synchronization (representing a disrupted functional network) and diminished gap junction coupling (representing a disrupted structural network).

The functional *β*-cell network has been suggested to follow a scale-free distribution^22^ with high clustering^24,25^, which implies a subset of highly connected nodes called “*β*-cell hubs”^18,23^. Importantly, *β*-cell hubs can exert a strong influence over islet dynamics^21,24,25^ and are preferentially disrupted in diabetes^23^. However, it is unclear whether the scale-free distribution in the functional network results from structural gap junction coupling or intrinsic properties of the *β*-cells themselves, such as the activity of Glucokinase (GK) or ATP-sensitive K^+^ (K_ATP_) channels which drive *β*-cell excitability.

Alternatively, proxies for assessing the structural network, such as electrophysiological measurements or computational studies that include electrical coupling, suggest it is not feasible for one cell or a small population of cells to have disproportionate influence over the islet dynamics^26,28–30^ and that specialized subpopulations are not required to achieve coordinated dynamics^31^. These seemingly contradicting results have caused debate over the most accurate topological description of the islet network^32,33^. We hypothesize these apparent contradictions can be explained through a careful study of the relationship between functional and structural networks.

Here, we use experimental and computational techniques to investigate the relationship between the islet functional and structural networks, with the intention of settling these highly debated questions relating to the role of *β*-cell heterogeneity. We frame our study around three questions. First, *are subpopulations that emerge from the β-cell functional network differentiated by a unique location in the structural network or by intrinsic properties of the cells?* While sometimes assumed, it is unclear whether the role of *β*-cell hubs in the functional islet network is due to a unique structural position or unique intrinsic dynamics. We hypothesize that this distinction may clarify whether *β*-cell hubs influence islet dynamics by dominating electrical communication. Further, the debates around islet heterogeneity and dynamics assume that the functional network directly reflects the structural network, which is not always true^17,20^. This leads to our second question: *what does the islet functional network indicate about its underlying structure or intrinsic cell properties on an individual cell basis?* Finally, as the robustness of *β*-cell networks change in diabetes, we investigated the islet functional network upon changes to the structural network. Specifically, *can we fully explain the disruption in the functional network by the removal of nodes and edges in the structural network, or are there other factors involved?* By answering these questions, we provide insight into the relationship between structure, function, and intrinsic cell dynamics in the pancreatic islet.

## Results

### Cellular metabolism, but not elevated gap junction coupling, is observed in highly synchronized cells in a simulated *β*-cell network

To investigate the relationships between functional and structural networks, we first analyzed a simulated *β*-cell network. We used a well-validated multi-cellular coupled ODE model, which describes the electrophysiological properties of the *β*-cells^34–37^ and includes heterogeneity in factors underlying cellular excitability and gap junction electrical coupling (**Figure 1a**).

**Figure 1:**
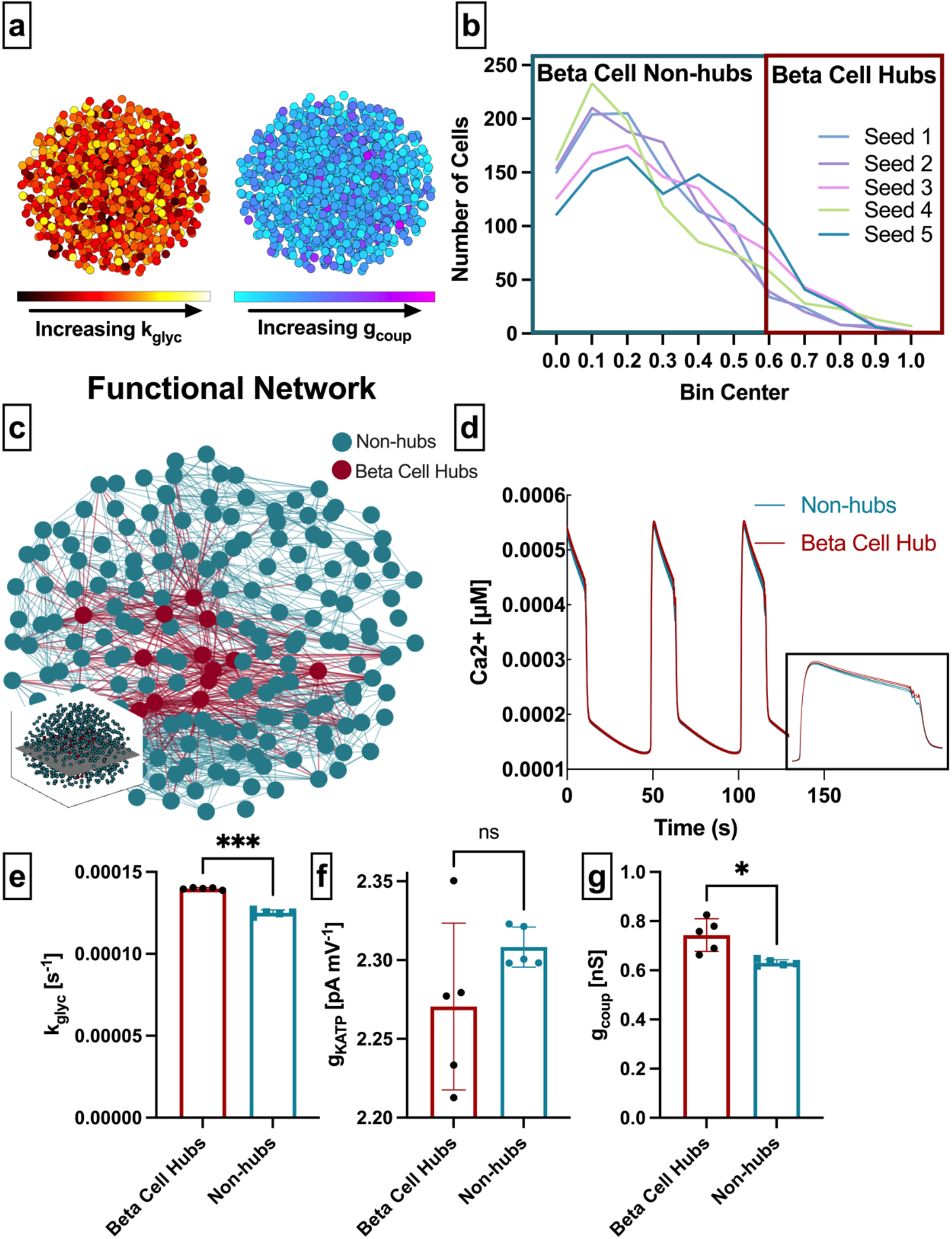
Analysis of parameters underlying highly connected cells in a simulated islet network. **a:** schematic of 1000 *β*-cell computational model, with cells false-colored by heterogeneity in k_glyc_ (left) and gcoup (right) parameter values. **b:** Distribution of functional connections (edges), determined from functional network analysis. Colors show five simulated islets. Hub cells (red outline) are any cell with > 60% of maximum number of links. **c:** 2-dimensional slice of the simulated islet, with lines (edges) representing functional connections between synchronized cells. Hub cells indicated in red. Slice is taken from middle of islet (see inset). **d:** Representative [Ca^2+^] time-course of a hub (red) and non-hub (blue). Inset shows zoomed enlarged signal from 45 – 80 seconds. **e:**Average rate of glucose metabolism (*k_glyc_*) values compared between hub and non-hub, retrospectively analyzed. **f:** as in e for maximum conductance of ATP sensitive potassium channel (g_KATP_). **g:** as in e for gap junction conductance (*g_coup_*). Significance in e-g was determined by paired Students t-test. \**P* ≤ 0.05 and \*\*\**P* ≤ 0.001.

We investigated our first question, whether subpopulations that emerge from the *β*-cell functional network differentiated by a unique location in the structural network or by intrinsic properties of the cells? We extracted the islet’s functional network by assigning a node to each cell and an edge between any cell pair whose temporal intracellular free-Ca^2+^ ([Ca^2+^]) traces had a correlation coefficient (*R_ij_*) greater than a threshold *R_th_*. The degree was defined as the number of edges a given node possesses (**Table 1**). We set the threshold to *R_th_* = 0.9995, such that the normalized degree distribution had many low degree cells and few high degree cells (**Figure 1b** and **Table 1**) to match prior studies^22,23^. We then identified *β*-cell hubs as cells whose normalized functional network degree was greater than 60%^23^ (**Figure 1b, c**).These *β*-cell hubs tended to depolarize before and repolarize after the islet average (**Figure 1d**), in agreement with previous studies^23,38^.

**Table 1:**
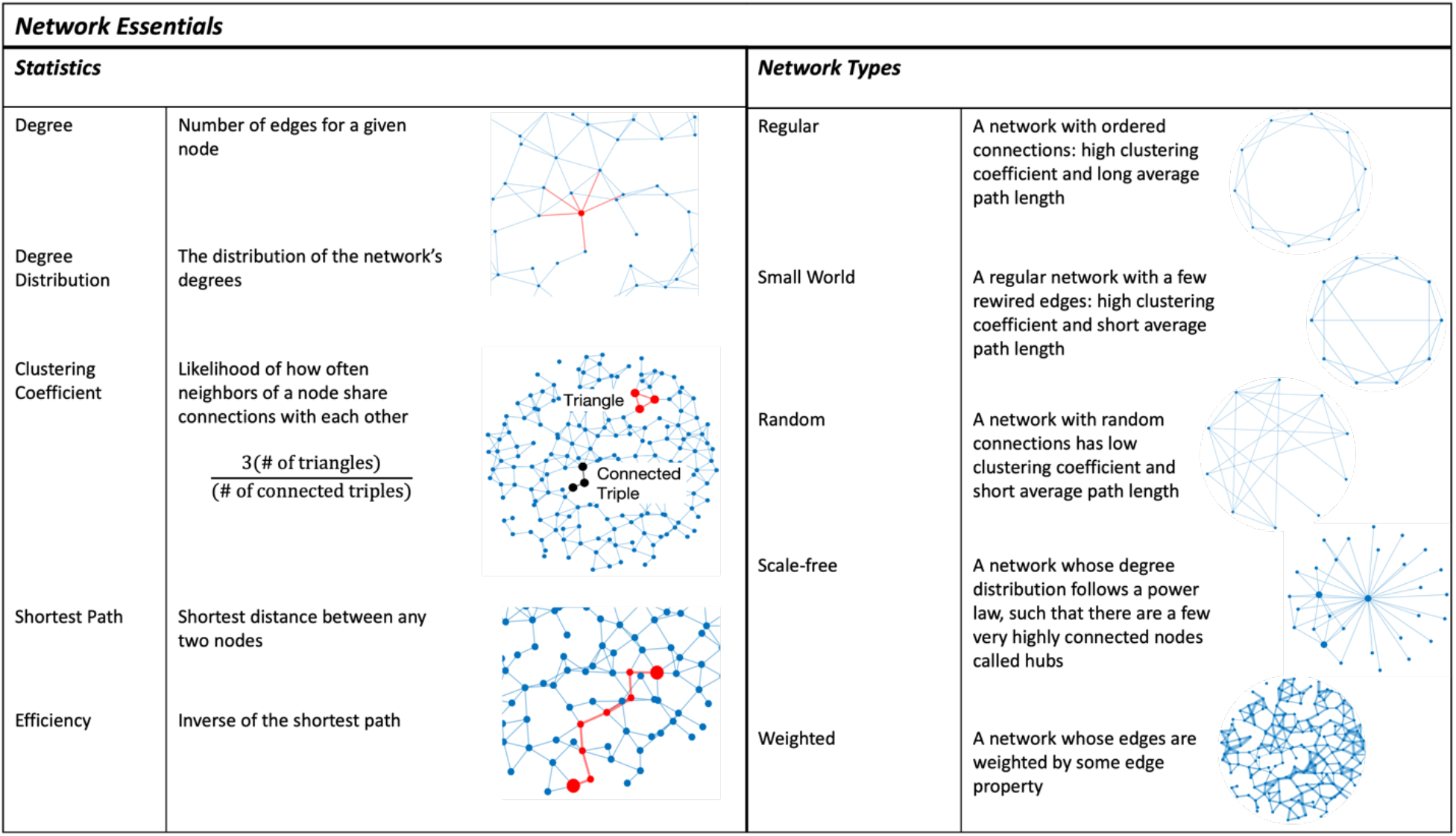
Essential network statistics and types. Left: Five network statistics used to quantify the network in this paper. Representative networks show an example of the statistic in red and the rest of the network in blue. Right: Five network types referred to in this paper. Regular, small world, and random networks are all made with the same number of nodes and edges, but different configurations of edges. Scale free network shows three “hub” nodes where node size is proportional to degree.

| Network Essentials |  |  |  |  |
| --- | --- | --- | --- | --- |
| Statistics |  |  | Network Types |  |
| Degree | Number of edges for a given node |  | Regular | A network with ordered connections: high clustering coefficient and long average path length |
| Degree Distribution | The distribution of the network's degrees |  | Small World | A regular network with a few rewired edges: high clustering coefficient and short average path length |
| Clustering Coefficient | Likelihood of how often neighbors of a node share connections with each other<br>$\frac{3(\text{\# of triangles})}{(\text{\# of connected triples})}$ | | Random | A network with random connections has low clustering coefficient and short average path length |
| Shortest Path | Shortest distance between any two nodes |  | Scale-free | A network whose degree distribution follows a power law, such that there are a few very highly connected nodes called hubs |
| Efficiency | Inverse of the shortest path |  | Weighted | A network whose edges are weighted by some edge property |

We then examined the relationship between functional network-derived *β*-cell hubs and the rate of glucose metabolism (*k_glyc_*) or maximum K_ATP_ conductance (g_KATP_). *β*-cell hubs had a significantly higher rate of glucose metabolism (*k_glyc_*) than non-hubs (*p* ≤ 0.0001)(**Figure 1e**) but were not different in their maximum K_ATP_ conductance (g_KATP_) (*p* = 0.18)(**Figure 1f**). These results indicate that *β*-cell hubs are differentiated by glucose metabolism rather than K_ATP_-dictated cellular excitability. Interestingly, *β*-cell hubs had only a slightly higher gap junction conductance (*g_coup_*) than non-hubs (*p* = 0.032)(**Figure 1g**). To ensure consistency in our findings, we repeated the analysis with various thresholds (**Supp. 1a**). For all thresholds analyzed, the difference between hubs and non-hubs was greater for *k_glyc_* than *g_coup_* (**Supp. 1b**), with *k_glyc_* being significantly higher in hubs for all five thresholds and *g_coup_* being significantly higher in hubs for only two thresholds. These results indicate that both metabolic activity and gap junction coupling correlate with high cellular synchronization, where metabolic activity has a stronger correlation.

### Elevated metabolic activity, but not elevated gap junction permeability, is observed experimentally in highly synchronized cells

We next investigated the subpopulations that emerge from the *β*-cell functional network experimentally. We performed time-lapse imaging of [Ca^2+^] within islets isolated from Mip-CreER;Rosa-LSL-GCamp6s mice that express GCamP6s in *β*-cells (*β*-GCamP6s mice, see methods). At elevated glucose, islet *β*-cells showed synchronous oscillations in [Ca^2+^] (**Figure 2a,b**). We extracted the functional network (**Figure 2c**) and again generated a normalized degree distribution (**Figure 2d**) which reflects a scale-free-like distribution.

**Figure 2:**
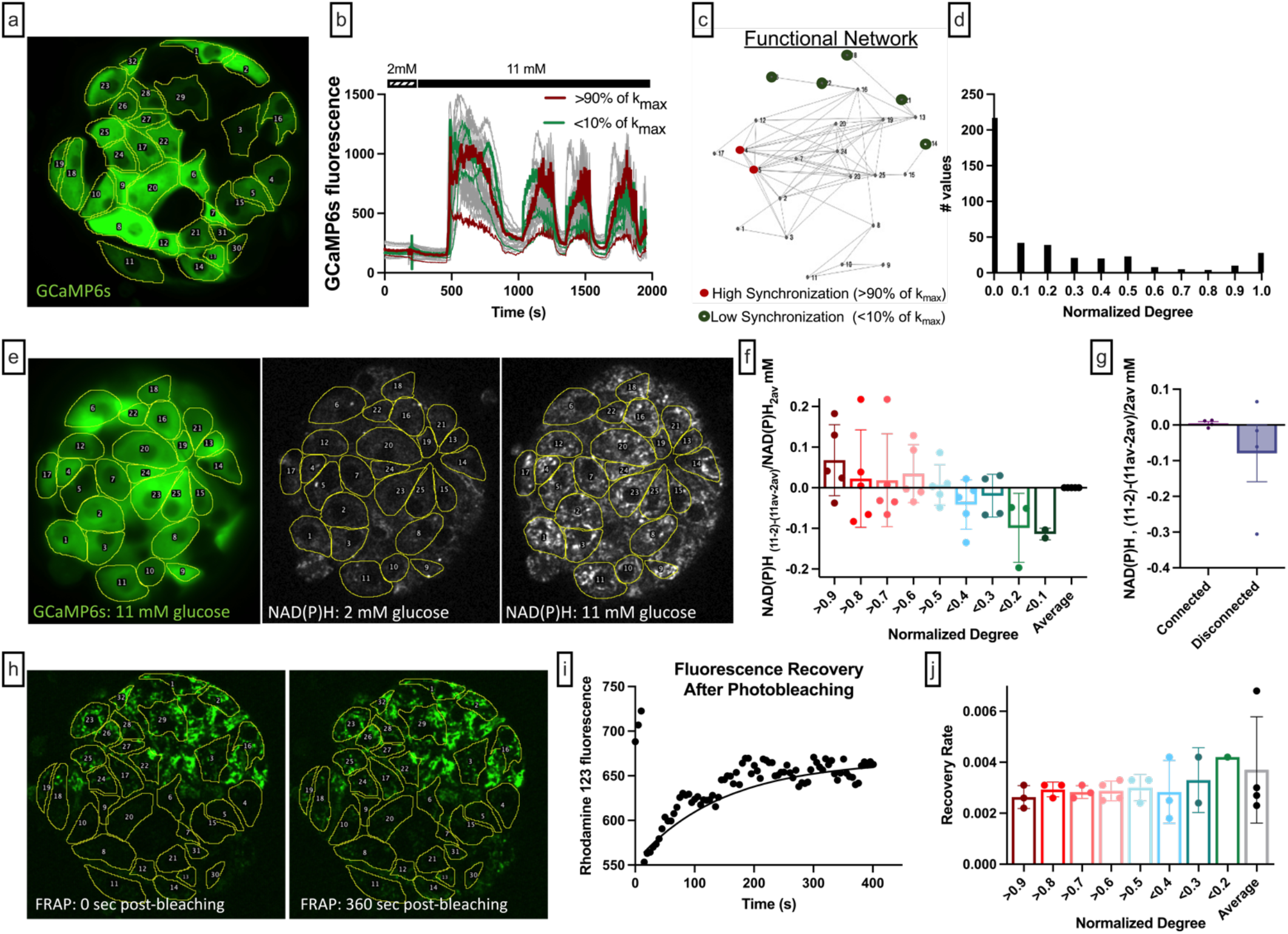
Comparison of the calcium-based network with the metabolic activity, NAD(P)H and coupling conductance. **a:** Mouse pancreatic islet expressing fluorescent calcium indicator GCaMP6s in *β*-cells. Glucose level 11 mM. **b:** GCamp6s time traces recorded at 2- and 11-mM glucose. Red curves represent dynamics of the most coordinated cells. These cells had highest number of edges, i.e., normalized degree >0.9. Green curves represent dynamics of the least coordinated cells, i.e., normalized degree <0.1, the rest of the cells are shown in grey. **c:** Ca^2+^ - based functional network of the islet shown in (e). Red dots represent most coordinated cells, and green dots – least coordinated cells which had at least 1 edge. **d:** Degree distribution of all eight analyzed islets. Threshold of R_th_=0.9 was used to filter the Pearson-based coordination matrix and obtain the functional network. **e:** *Left:* Mouse pancreatic islet expressing fluorescent calcium indicator GCaMP6s. Glucose level 11 mM. *Middle:* NAD(P)H autofluorescence of the islet and the same cell layer, recorded at 2 mM glucose. *Right:* NAD(P)H autofluorescence recorded at 11 mM glucose. **f:**Change of the NAD(P)H levels in each cell in response to glucose elevation, with respect to the islet-average change. Metabolic activity here is compared for connected and disconnected cells. **g:** Change in NAD(P)H levels in response to glucose elevation, with respect to the islet-average change: here comparison is made for the most coordinated cells (normalized degree >0.9) with the less coordinated cells (normalized degree >0.8, >0.7, >0.6, … <0.2, <0.1). f,g represent n=5 islets with a total of 131 cells. **h:** Rhodamine123 fluorescence of the islet and the same cell layer as in **a**, recorded immediately after photobleaching of the bottom half of each islet (**left**) and recorded at 360 seconds after the photobleaching (**right**). **i:** Time trace of the fluorescence recovery after photobleaching (FRAP) of the lower half of each islet, shown in **h**. The rate of recovery is proportional to the Cx36 gap junction permeability of each cell. **j:** FRAP recovery rate (sec^−1^) in each of the photobleached cells: here the comparison is made for the most coordinated cells (normalized degree >90%) with the less coordinated cells (normalized degree >80, >70, >60,…<20, <10%). j shows results from 4 islets and 202 cells). All the cells in this graph had a degree larger than 0.01.

To test whether cellular metabolic activity differentiated high-degree cells from normal or low-degree cells, we measured two-photon excited NAD(P)H autofluorescence in conjunction with tim-elapse imaging of [Ca^2+^] dynamics at 2mM and 11mM glucose (**Figure 2e**). The metabolic response of individual *β*-cells in the islet was calculated as the de-meaned difference between NAD(P)H autofluorescence at high glucose compared to low glucose (*NAD(P)H*_11-2_-*NAD(P)H_11av-2av_*). This metric was 0 if the NAD(P)H response of a cell was similar to the islet average, >0 if a cell showed an elevated metabolic response, and <0 if a cell showed a reduced metabolic response. The NAD(P)H response decreased as the cell’s normalized degree decreased (**Figure 2f**). This indicates that highly synchronized cells are more metabolically active, in agreement with findings from simulated islets. Furthermore, the NAD(P)H response trended lower in cells that were functionally disconnected (Ca^2+^ oscillations lacking any synchronization), compared to connected cells (**Figure 2g**). Therefore, the functional network is strongly related to cellular glucose metabolism which drives islet dynamics.

Fluorescence recovery after photobleaching (FRAP) measurements of dye transfer kinetics can quantify gap junction permeability^39^. We performed FRAP measurements in conjunction with time-lapse imaging of [Ca^2+^] dynamics, to map cellular gap junction connections in the same cell layer as the functional network (**Figure 2h,i**). The rate of recovery, which assesses gap junction permeability, lacked any clear trend with [Ca^2+^] synchronization **(Figure 2j**). This indicates that cell synchronization is not correlated with gap junction permeability, as measured by FRAP, consistent with findings from simulated islets. *Therefore, the functional network is not strongly related to the structural network*.

### Synchronization between a cell pair more likely indicates shared metabolic properties than shared gap junction coupling

We next investigated our second question: what does the islet functional network indicate about its underlying structure or intrinsic properties on an individual cell basis? Irrespective of a cell’s functional network degree, it was rare for a cell pair to be connected in both the functional and structural networks (**Figure 3a**). The probability that two cells were synchronized in the functional network, given that they shared a gap junction in the structural network, was *Pr*(*Sync|GJ*) = 0.39 (**Figure 3b**). Therefore, the presence of a structural edge does not imply a functional edge. The alternative probability of a structural edge given a functional edge was *Pr*(*GJ|Sync*) = 0.04 (**Figure 3b**), indicating that very few functionally coupled cells are connected by gap junctions. These findings indicate that structurally connected cells and functionally connected cells are two distinct groups with little overlap (*Pr*(*both*) = 0.2) (**Figure 3c**).

**Figure 3:**
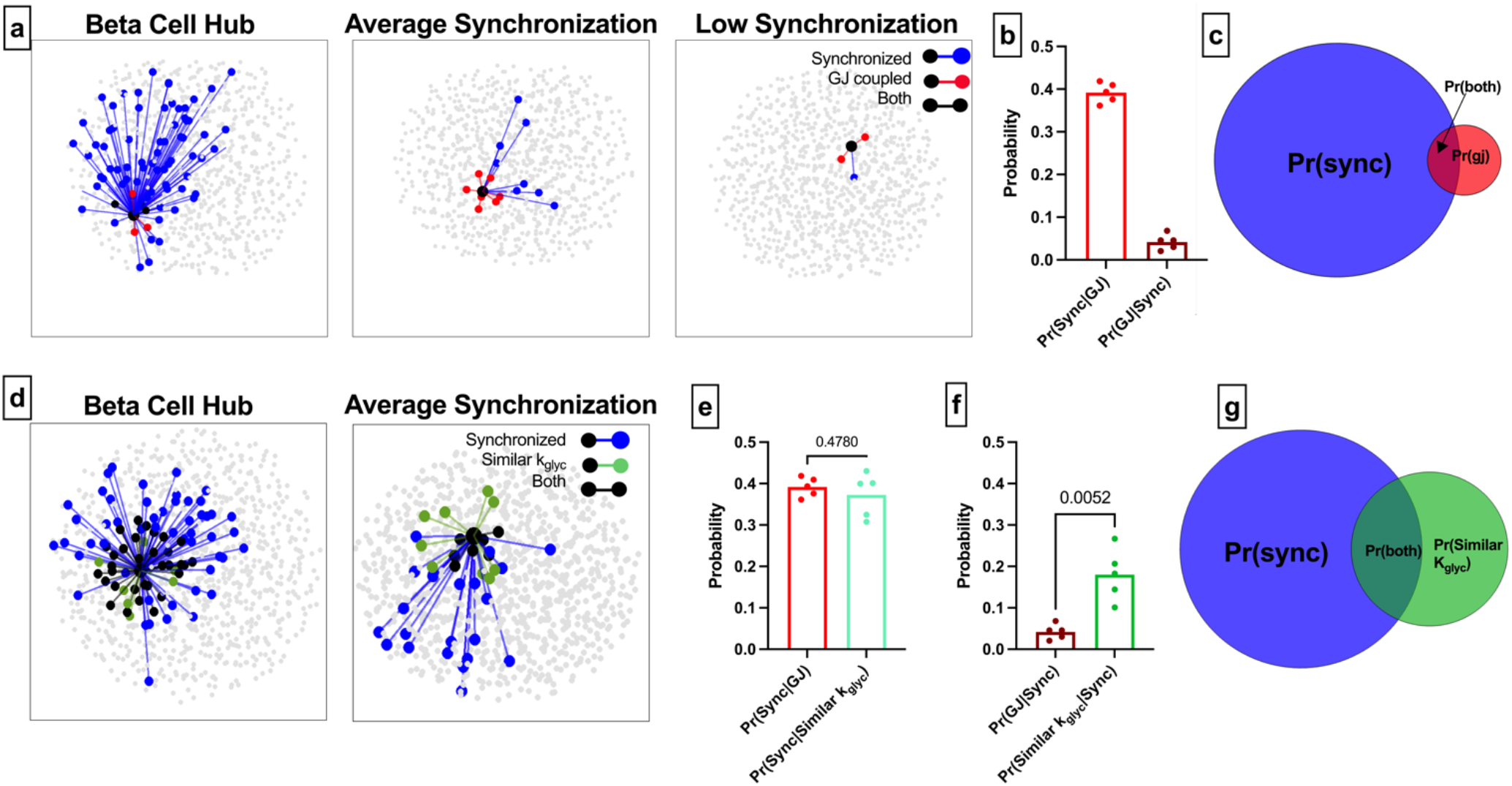
Comparison of functional connections, gap junction junctions, and glucose metabolism. **a:** Functional connections and structural connections for a representative highly connected *β*-cell hub, average cell, and low connected cell. Blue cells and edges indicate cells determined as “synchronized” via functional network analysis. Red cells are GJ coupled. Black cells are both GJ coupled and synchronized. **b:** Probability that a cell is synchronized given it is gap junction coupled, or vice versa, that a cell is gap junction coupled given it is synchronized. **c:** Ven diagram showing overlap between the synchronized cells and gap junction coupled cells within the simulated islet. **d:** Functional connections and metabolic connections for a *β*-cell hub and a cell with averaged synchronization. Blue cells and edges indicate cells determined as “synchronized” via functional network analysis. Green cells and edges indicate cells that have similar k_glyc_ and are within 15% islet from each other. Black cells and edges indicate both synchronized and similar k_glyc_. **e:** Probability that a cell pair is synchronized given it shares a gap junction, has high average kglyc or has similar k_glyc_. **f:** Probability that the cell pair is gap junction coupled, has higher than average k_glyc_ and within 15% of islet distance (*Pr* = 0.0050), or has similar k_glyc_ and within 15% of islet distance (*Pr* = 0.0052), respectively, given the pair is synchronized. **g:** Ven diagram showing overlap between the synchronized cells and metabolically similar cells. Shaded area in c and g is proportional to indicated probability. Significance in e and f was determined by a repeated measures one-way ANOVA with Tukey’s multiple comparisons \*\**P* ≤ 0.01.

We analyzed the sensitivity of these measures to the synchronization threshold *R_th_*. Tautologically, as the threshold was increased, the number of synchronized cells decreased (**Supp. 1d**), causing *Pr*(*GJ|Sync*) to increase (**Supp. 1e**). However, the “overlap” between the functional network and structural network did not increase (**Supp. 1f**). Our findings that the functional and structural networks are two distinct groups with little overlap is consistent across thresholds.

These data corroborate with initial findings showing that function is not strongly correlated with structure. To statistically investigate whether metabolic properties are correlated with function, as shown in our previous results, we created a new network where edges were drawn between cells with similar rates of glucose metabolism (k_glyc_) (see methods) **(Figure 3d**). Parameters were chosen such that there was no difference between *Pr*(*Sync|Similar k_glyc_*) and *Pr*(*Sync|GJ*) (**Figure 3e**), allowing for a direct comparison between probabilities without a strong influence of sample size. The alternative probability *Pr*(*Similar k_glyc_|Sync*) was significantly higher than *Pr*(*GJ|Sync*) (**Figure 3f**). If two cells have synchronized Ca^2+^ oscillations, they are more likely to contain unique metabolic characteristics than be gap junction coupled. This is further indicated by the increased overlap between the [Ca^2+^] synchronization-derived functional network and the metabolic activity-derived network (**Figure 3g**). These results further indicate that metabolic activity, not gap junction connections, is a greater driving factor for cells to show high [Ca^2+^] synchronization and thus influence the functional network.

### Elevated glucose metabolism is a greater driver of long-range functional connections

In accordance with the islet cytoarchitecture, gap junction connections only exist between highly proximal cells. However, there is no distance constraint on [Ca^2+^] oscillation synchronization (**Figure 4a**). To determine whether this spatial constraint was responsible for the low correspondence between the functional and structural networks, we asked whether a series of highly conductive gap junction connections could predict long-range functional connections. We calculated the path which maximized conductance (see methods: shortest weighted path length) after weighting the structural network by g_coup_ (**Figure 4b**). Not all synchronized cell pairs had chains of larger total g_coup_ than non-synchronized cell pairs (**Figure 4b**). Further, the probability distributions of total g_coup_ for synchronized cells highly overlapped with those for non-synchronized cells, irrespective of distance (**Figure 4c,d, and Supp. 2a**). However, the total conductance normalized by separation distance was significantly less for non-synchronized cells than for synchronized cells (**Figure 4e**). Thus, long-range functional connections traverse cells that are connected on average by slightly higher gap junction conductance. Therefore, while gap junction conductance can influence long-range synchronization, the substantial overlap between synchronized and non-synchronized cells indicates that it is not the driving factor and cannot be used to differentiate cell pairs connected in the functional network.

**Figure 4:**
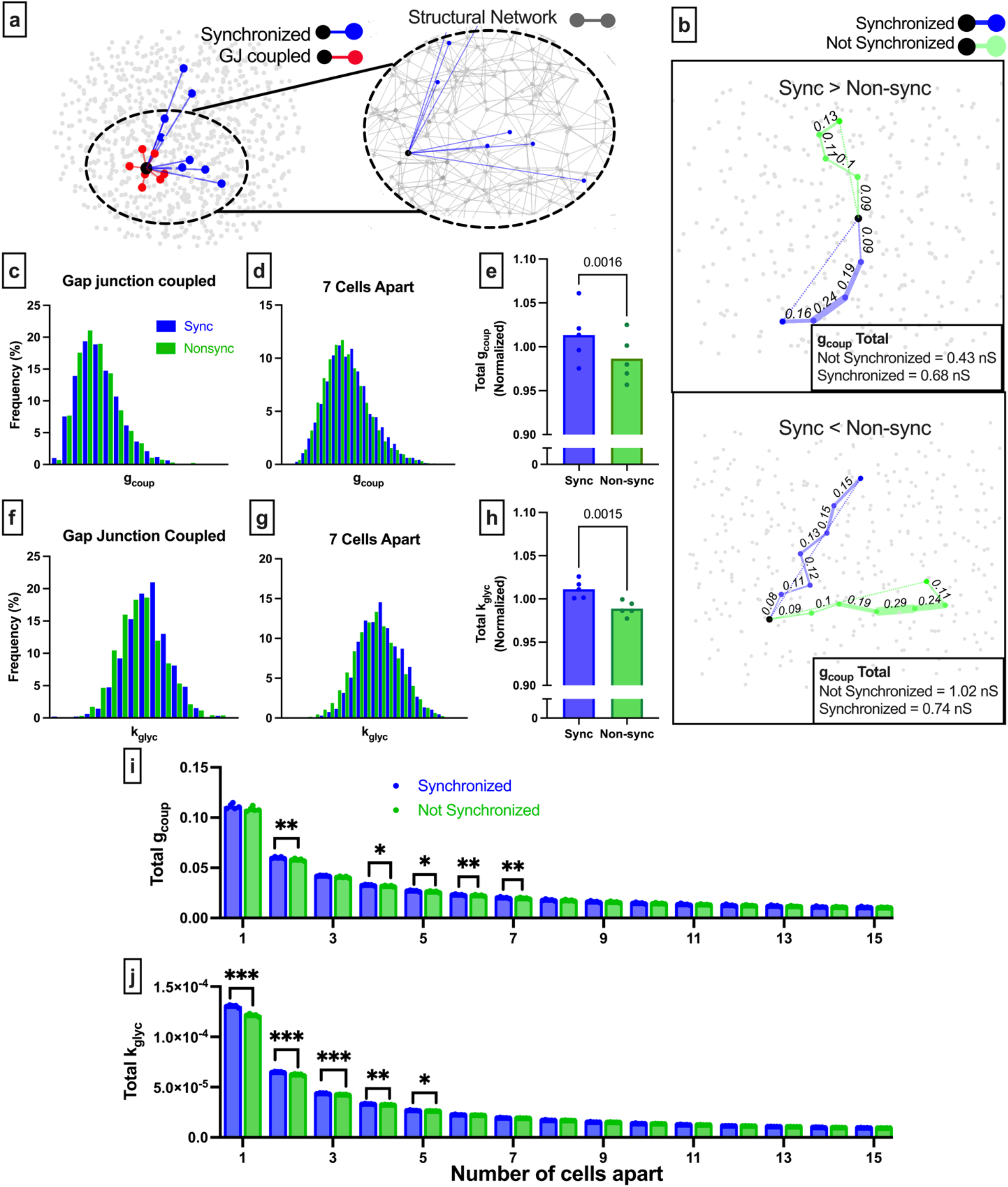
Comparison of long-range functional synchronization, gap junction network, and glucose metabolism. **a:** Functional connections and gap junction connections for a representative cell (black). Synchronized cells in blue, gap junction coupled cells in red. Inset shows the entire structural network (grey) with functional connections for the same cell shown in blue. **b:** Two representative cells (black) and the shortest path to a synchronized cell (blue) and a non-synchronized cell (green). Path is weighted (shown by edge thickness) by g_coup_. For the cell on the top panel, the synchronized path has a higher cumulative gap junction conductivity (0.68 nS) than the nonsynchronized path (0.43 nS). For the cell on the bottom panel, the synchronized path has a lower cumulative gap junction conductance (0.74 nS) than the nonsynchronized path (1.02 nS). **c:** Probability distribution of total gcoup for synchronized cells (blue) and non-synchronized cells (green) that are directly connected by gap junctions. **d:** As in c for cells pairs that are 7 cells apart. **e:** Comparison of the total g_coup_, normalized by distance for synchronized cells (blue) and non-synchronized cells (green). Each dot indicates the average resistance for a single simulated islet. **f:** As in c but with connections weighted by rate of metabolism. **g:** As in d but with connections weighted by the rate of glucose metabolism. **h:** As in e but with connections weighted by the rate of glucose metabolism. **i:** Total g_coup_ for synchronized and nonsynchronized cells organized by cell distance. **j:** As in i but for the metabolically weighted network. Significance in e,h was assessed by a two-tailed paired t-test, with P value indicated. Significance in i and j was, assessed by paired t-tests with Bonferroni correction. Reported P values are adjusted. \**P* ≤ 0.05, \*\**P* ≤ 0.01, \*\*\**P* ≤ 0.001.

To examine how the cellular glucose metabolism influences long-range functional connections, we repeated this procedure but weighted graph edges by *k*_glyc_ rather than g_coup_. Again, the probability distributions of the total *k*_glyc_ for synchronized cells were highly overlapping with that for non-synchronized cells; irrespective of distance (**Figure 4f,g** and **S3**). The total rate of glucose metabolism, normalized by the separation distance, was significantly less for non-synchronized cells than for synchronized cells (**Figure 4h**). Thus, on average, long-range functional connections traverse cells with higher glucose metabolism.

To test whether there was a spatial relationship for glucose metabolism or gap junction conductance-controlled synchronization, we analyzed the total conductance or total rate of glucose metabolism for different separation distances and between synchronized or non-synchronized cell pairs. Total g_coup_ was significantly higher for synchronized cells 2 to 7 cells apart **(Figure 4i**). Total *k*_glyc_ was significantly higher for synchronized cells 1 to 5 cells apart, with the significance larger than that of total g_coup_. Thus, cell pairs with similar *k*_glyc_ can strongly influence function connections up to 5 cells apart, while high g_coup_ can influence function connections over slightly longer distances (5-7 cells apart).

### Individual gap junction connections cannot be predicted by the functional network when gap junction conductance correlates with metabolic activity

Prior work suggested that cells with highly synchronized Ca^2+^ oscillations possess both elevated metabolic activity (in agreement with our observations, **Figure 2**) *and* high levels of gap junction conductance (unlike what we observe experimentally, **Figure 2**)^23,27^. To address this possibility, we repeated our analyses on a simulated islet in which the rate of glucose metabolism positively correlated with gap junction conductance (**Supp. 4**). Our results were consistent across correlated and non-correlated distributions.

### Modulating the islet-wide structural network in C×36 knockout islets

Our third question - can we fully explain the disruption in the functional network by the removal of edges in the structural network - is particularly relevant to pathophysiological conditions, where gap junction coupling is disrupted. Reducing C×36 gap junction coupling will reduce electrical communication between *β*-cells, and thus removes edges from the structural network. We performed functional network analysis on islets from wild-type mice (C×36^+/+^), heterozygous C×36 knockout mice with ~50% reduced gap junction coupling (C×36^+/−^), and homozygous Cx36 knockout mice with no gap junction coupling (C×36^+/-^)^1^ (**Figure 5a-c**). We used a single synchronization threshold *R_th_* across all genotypes such that the degree distribution of WT islets showed a scale-free-like distribution^22^ and had an average of between 5-15 connections (see methods: Threshold).

**Figure 5:**
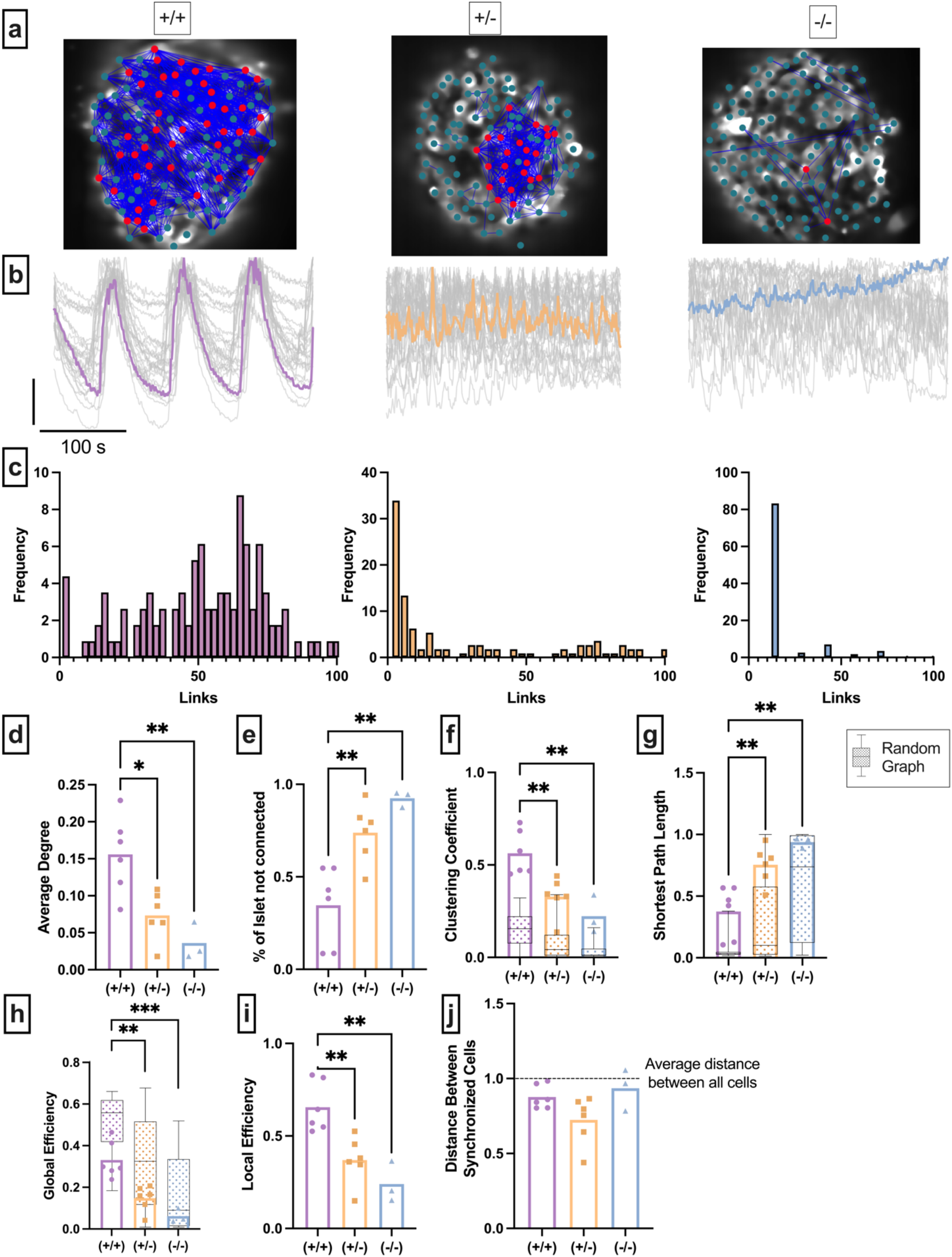
Experimental islet functional network is influenced by changes in the structural network. **a:** Representative images of C×36^+/+^ islet (left), heterozygous C×36^+/-^ islet (middle), and homozygous islet (right), overlaid with a synchronization network. Dot signifies a cell, blue lines represent functional network edges that connect synchronized cells. Red cells are *β*-cell hubs. **b:** Oscillations in [Ca^2+^] of corresponding islet in a. grey lines represent time-course for each cell, colored line represents mean islet time-course. **c:** Frequency histograms of connections for the corresponding islet in *a*. Connections are calculated using functional network analysis and normalized to the maximum degree. As such 100% links represent the cell with the greatest total number of connections. **d:** Normalized degree of functional network between cell pairs averaged over islets of each C×36^+/+^, C×36^+/-^, C×36^-/-^ mouse. **e:** Percent of cells in the islet which have are not connected to any other cell (eg. All *R_ij_* < *R_th_*). **f:** Average clustering coefficient. Overlaid is the box and whisker plot of 1000 random networks created with the same number of nodes, edges, and degree as the islet functional network. **g:** Average Shortest path length. Overlaid is the box and whisker plot of 1000 random networks per islet, as in f. **h:** Average Global Efficiency. Overlaid is the box and whisker plot of 1000 random networks per islet, as in f **i:** Local efficiency. **j:** Average number of pixels between synchronized cells normalized to the average number of pixels between all cells in the islet. *D* < 1 indicates that cells preferentially synchronize to nearby cells. Significance in d-k determined by ordinary one-way ANOVA with Tukey multiple comparison test. Reported P values are adjusted. \**P* ≤ 0.05, \*\**P* ≤ 0.01, \*\*\**p* ≤ 0.001.

**Figure 5c** shows representative degree distributions of functional networks corresponding to the levels of C×36, with average degree distributions for all islets in **Supp. 5a**. Results separated by mice are shown in **Supp. 6.** C×36^+/−^ islets demonstrated decreased average network degree compared to wild-type C×36^+/+^ islets (**Figure 5d**), and homozygous C×36^-/-^ demonstrated further decreased degree compared to both wild-type C×36^+/+^ islets (**Figure 5d**) and C×36^+/-^ islets (**Figure 5d**), indicating that decreases in gap junction conductance lead to decreases in overall synchronization. Consistent with this, more cells in C×36^+/-^ islets (74% of cells) and C×36^+/-^ islets (93% of cells) were not functionally connected to any other cell, compared to in Cx36^+/+^ islets (40% of cells) (**Figure 5e**). These data show the functional network becomes highly sparse as edges in the structural network decrease. Islets were of similar size ranges in across each genotype (**Supp. 4c**).

We next sought to quantify how the removal of edges in the structural network influences the functional network topology (**Table 1**). Network topology statistics can provide insight into how the network functions. *Clustering coefficient* (*C_avg_*) quantifies the percent of nodes connected to a given node that are also connected with each other (**Table 1**). A high clustering coefficient indicates that cells tend to synchronize with other cells that share some property. The functional network clustering coefficient decreased significantly as gap junction coupling decreased (**Figure 5f**). To statistically quantify whether the functional network topology was a result of random edge generation, we compared each metric with that determined from 1000 erdos-renyi random networks^16,19,22^ (**Table 1**). If gap junctions are the property that gives the functional network a high clustering coefficient, then reducing gap junction coupling should result in a clustering coefficient that is completely explained by the random graph. For all levels of Cx36, the clustering coefficient of the islet functional network was greater than that of a random network (**Figure 6f**, box and whisker plot). Similar to our previous findings (**Figure 4**) this further suggests that something other than the structural network contributes the functional network topology.

**Figure 6:**
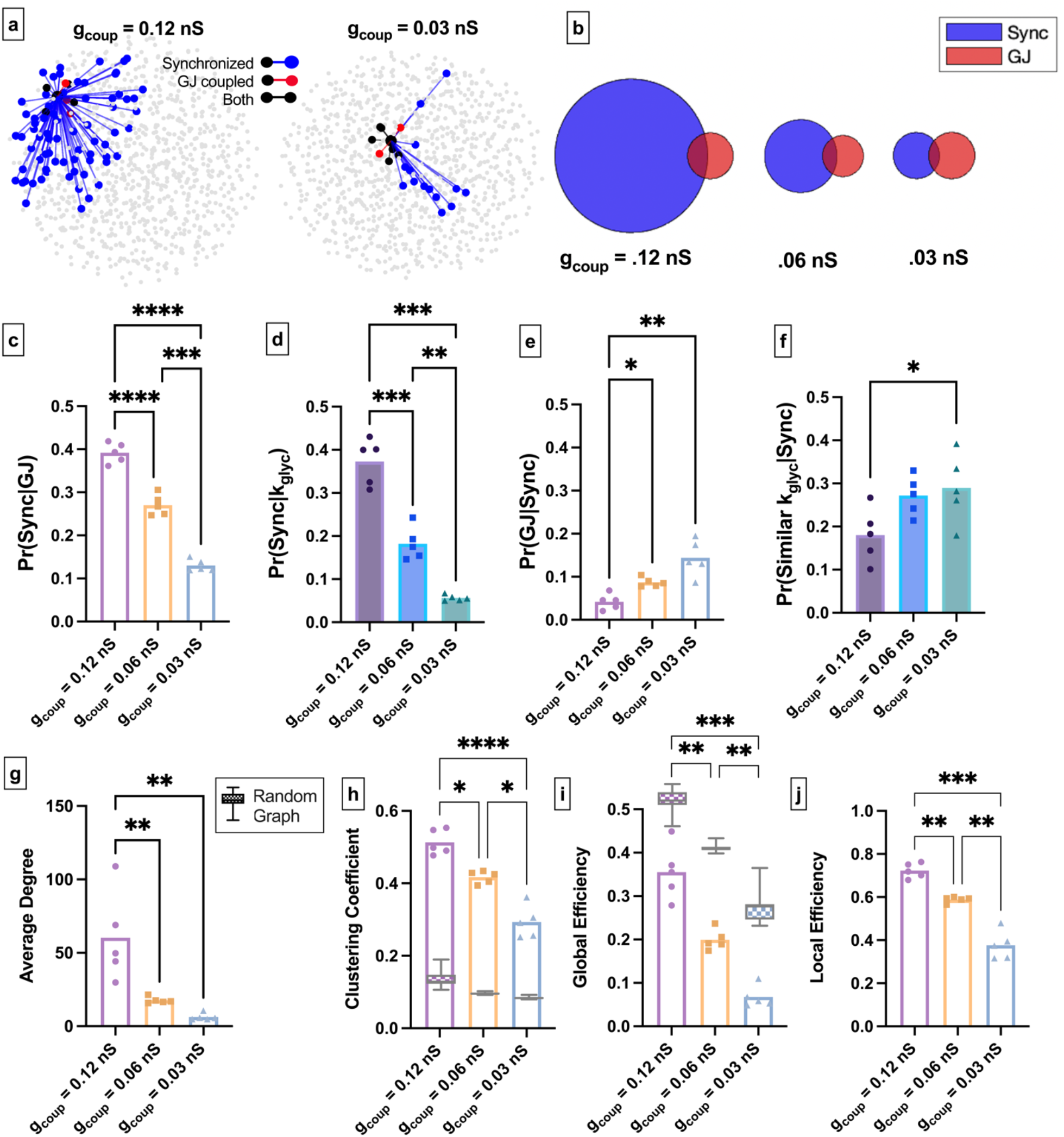
Simulated islet functional network is influenced by changes in the structural network. **a:** Representative cells within a simulated islet with high gap junction conductance (*g_coup_* = 0.12*nS*) (left) and within an islet with gap junction low conductance (*g_coup_* = 0.03 *nS*)(right). Connections between a representative cell are shown with synchronized cell pairs in blue, gap junction cell pairs in red, and both in black. All data points represent the average statistic for each mouse. **b:** Venn diagrams of the probability of synchronization and probability of gap junction with the probability of both synchronization and gap junction as the overlapped area. **c:** Probability that two cells are synchronized given they share a gap junction connection. **d:** Probability that two cells were synchronized given they had similar rate of glucose metabolism and were within 1islet the islet from each other. **e:** Probability that two cells shared a gap junction connection given they were synchronized. **f:** Probability of similar rate of glucose metabolism given two cells were synchronized. **g:** Global efficiency for 0.12*nS*, 0.06*nS*, and 0.03*nS*, for each islet seed. Overlaid is the box and whisker plot of 1000 random networks per simulated islet seed. **h:** Clustering coefficient for 0.l2nS, 0.06nS, and 0.03nS, for each islet seed. Overlaid is the box and whisker plot of 1000 random networks per simulated islet seed, as in g. **i:** Small world-ness of a network calculated using global efficiency. 1000 random networks per islet were created to determine uncertainty of each data point (shown in black). **j:** Local efficiency for 0.12*nS*, 0.06*nS*, and 0.03*nS*, for each islet seed. Shaded area in b is proportional to indicated probability. Statistical significance in c-j assessed by an ordinary one-way ANOVA with Tukey multiple comparison test. Reported P values are adjusted. \**P* ≤ 0.05, \*\**P* ≤ 0.01, \*\*\**P* ≤ 0.001.

The *average shortest path length* (*L_avg_*) is defined as the average smallest number of nodes required to traverse between any cell pair (**Table 1**). A random network or a regular network (see **Table 1**) with a few edges randomly moved will have a small shortest path length^16^. The normalized shortest path length increased as gap junction coupling decreased (**Figure 6g**). For C×36^+/+^ islets, the measured shortest path length was greater than that of a random network. As gap junction coupling decreased, the shortest path length showed a progressively larger intersection with that of a random network. For C×36^-/-^ islets all measurements could be explained by random chance. This suggests that the gap junction structural network can contributes to long-range properties of the functional network topology, similar to our previous findings (**Figure 4**). *Global efficiency*, which has been used to describe islet networks^22^, is related to the inverse shortest path length (**Table 1**). Global efficiency significantly decreased with decreased gap junction coupling, and again showed a larger intersection with that of a random network (**Figure 6h**). In all cases, these functional network characteristics did not show any significant mouse-to-mouse variability (**Supp. 6**).

The functional network in WT islets can exhibit *small-world qualities*^22^, meaning that it exhibits both high clustering of a regular network and high efficiency of a random network (**Table 1**). A small-word functional network is efficient to traverse (along functional edges), but cells tend to synchronize will other cells of similar attributes. We investigated whether small-world qualities were diminished when structural edges were removed. Previously^22^, small-worldness was quantified using random networks. However, as the functional network becomes sparser, the uncertainty in the random network-based approach increases (**Supp. 4d**, seen more clearly in **Supp. 6d**). To remove this uncertainty, we used an alternative metric^40^ which was formulated specifically to analyze small-worldness in sparse networks. *Local efficiency* reveals the efficiency of communication between neighbors of a node, when that node is removed. If both global efficiency *E_global_* and local efficiency *E_local_* are large, the network has small-world qualities^40^. Local efficiency significantly decreased with reduced gap junction coupling (**Figure 6i**).The decrease in both global and local efficiency with decreased gap junction coupling correspond to a decrease in the small world-ness of the functional network. Therefore, reducing gap junction coupling decreases (but does not completely abolish) the communication efficiency of the islet. However, the fact that random networks cannot explain the clustering observed with reduced gap junction coupling further indicates that other attributes are driving cellular synchronization.

### Simulating modulations to the gap junction structural network in the computational model

Finally, we combined all three of our questions by investigating how changes in the structural network affect the functional network topology and the relationships between function, structure, and cell glucose metabolism. We decreased structural edges by altering the average coupling conductance (*g_coup_*) in our simulated islet. As expected, islets with reduced average *g_coup_* showed reduced [Ca^2+^] oscillation synchronization: simulated islets with average *g_coup_* = 0.12*nS* had higher synchronization than those with average *g_coup_* = 0.06*nS* or *g_coup_* = 0.03*nS* (**Figure 6a**). Similarly, the total number of functional network edges decreased as *g_coup_* decreased (**Figure 6a,b**). *Pr*(*Sync|GJ*) decreased significantly as gap junction coupling was decreased (**Figure 6c**). This indicates that as the gap junction conductance decreases, the probability that two structurally connected cells will also be functionally connected decreases. Similarly, Pr(*Sync*|*k_glyc_*) decreased as gap junction coupling decreased (**Figure 6d**). This indicates that gap junctions play a role in synchronizing metabolically similar cells. Conversely, *Pr*(*GJ|Sync*) increased as gap junction coupling decreased (**Figure 6e**). The probability of two cells showing similar rates of glucose metabolism given that two cells were connected in the functional network Pr(*k_glyc_*|*Sync*) also increased as gap junction coupling decreased (**Figure 6f**). and showed higher probabilities than that for *Pr*(*GJ*|*Sync*). These results indicate that as gap junction coupling decreases, a synchronized cell is more likely to be connected via a gap junction or be metabolically similar. However, this is not because the overlap between the two networks is larger but because the number of synchronized cells is smaller (**Figure 6b**). Thus, given sufficient gap junction coupling, glucose metabolism drives synchronization between individual cells.

We then quantified the three-dimensional stimulated functional network topology upon reduced gap junction conductance and compared these with network topology calculated from the two-dimensional [Ca^2+^] imaging data. The average degree significantly decreased as coupling decreased (**Figure 6g**). The clustering coefficient *C_avg_* progressively decreased with decreasing gap junction conductance and was always substantially greater than that determined from a random network (**Figure 6e**). Global efficiency also progressively decreased with decreased gap junction conductance (**Figure 6i**) and was always less than that determined from a random network (**Figure 6i**). These trends are similar to experimental observations. The more pronounced lack of overlap between random networks may be due to having more cells from a three-dimensional islet network. The local efficiency decreased with gap junction conductance (**Figure 6j**), also similar to experimental observations. The decreases in both global efficiency and local efficiency with decreasing gap junction conductance indicate that the network strays away from small-world properties as gap junction conductance decreases.

## Discussion

Diabetes Mellitus is marked by changes in both the gap junction-based structural^1,12,41–44^ and [Ca^2+^] synchronization-based functional^11,12,45,46^ networks in the pancreatic islet. However, the relationship between these two network representations is not well understood and has caused confusion in our understanding of islet communication^26,29,31–33^. Our aim was to investigate the relationship between functional networks, structural networks, and underlying *β*-cell dynamics in the pancreatic islets. We framed our study around three guiding questions. 1) Are subpopulations that emerge from the *β*-cell functional network differentiated by a unique location in the structural network or by intrinsic properties of the cells? 2) What does the islet functional network indicate about its underlying structure or intrinsic cell properties on an individual cell basis? 3) Can we fully explain the disruption in the functional network by the removal of nodes and edges in the structural network or are there other factors involved? **Figure 7** shows a graphical representation of the answers to these questions.

**Figure 7:**
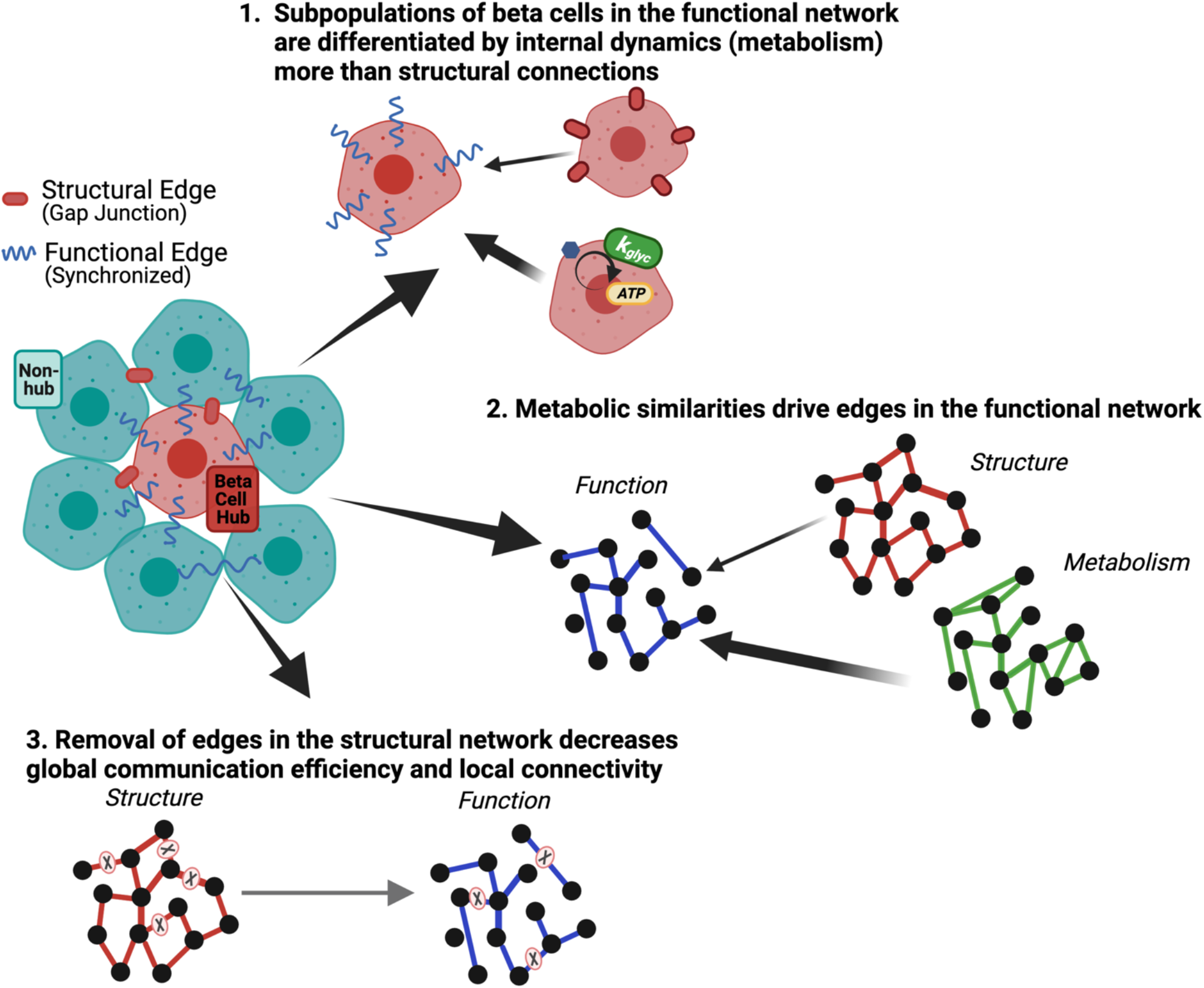
Graphical summary of Results. Graphical summary of results from data. To answer question 1: we show that sub populations of *β*-cells in the functional network are driven by metabolism more than gap junction coupling (Top). To answer question 2: we show that metabolic similarities drive edges in the functional network (right). To answer question 3: we show that removal of edges in the structural network decreased global communication efficiency and local connectivity (bottom).

In answering question 1, we concluded that subpopulations of *β*-cells in the functional network are driven by metabolic activity more than gap junctions. Our computational and experimental results show a subpopulation of highly synchronized cells, known as *β*-cell hubs^23,47^ (**Figure 1**). *β*-cell hubs were strongly associated with elevated metabolic activity; but associated only weakly (by computational model, **Figure 1**) or not at all (by experiment, **Figure 2**) with structural gap junction coupling. Semi-quantitative immunofluorescence previously demonstrated elevated glucokinase protein in *β*-cell hubs, suggesting elevated metabolic activity^23^. However, evidence for elevated gap junction coupling in *β*-cell hubs was based on [Ca^2+^] synchronization analysis, in disagreement with our finding here. *Thus, gap junctions are not the primary distinguishing factor in determining variations in cellular synchronization*. Computational studies have shown that the hyperpolarization of *β*-cell hubs disproportionately disrupts islet [Ca^2+^] dynamics, but the actual removal of hubs does not^26,28^. The conclusion that *β*-cell hubs are uniquely defined by metabolic activity rather than gap junction coupling, helps explain these results. If *β*-cell hubs drive synchronization via elevated metabolic activity, hyperpolarizing them excessively decreases average metabolism across the islet, thus disproportionately affecting excitability while removing them allows the next most metabolically active cells to fill the hub cell role (as described in^28^). Corroborating this hypothesis, prior experiments have shown that cells that are preferentially activated by optogenetic ChR2 stimulation were more metabolically active by NAD(P)H autofluorescence, but not preferentially gap junction coupled^37^.

In answering question 2, we concluded that the functional network reflects variations in cellular metabolic activity. Conditional probabilities quantify the relationship between structural and functional networks at an individual cell basis. The probability of high synchronization given gap junction connections (Pr(Sync|GJ)) was only ~40% (**Figure 3b**). The fact that a minority of structurally connected cells were functionally connected, indicates that additional factors contribute to the synchronization of cell pairs. We found the probability of a gap junction connection given a synchronized cell pair (Pr(GJ|Sync)) was <5%, indicating that high [Ca^2+^] synchronization rarely implies a gap junction connection (**Figure 3b**). However, the probability that two synchronized cells had similar metabolic activity or elevated metabolic activity was significantly larger (~20%) (**Figure 3e,f**). *Therefore, the functional network reflects variations in cellular metabolic activity rather than structural gap junction connections*. Over long-range functionally connected cell pairs, we observed similar results (**Figure 4**). While synchronized cells had a slightly higher gap junction conductance than non-synchronized cells, the strong overlap in distributions indicates that long-range functionally connected cell pairs are still not always determined by a path of elevated gap junction conductance. The surprising part of these findings is that gap junctions do not guarantee high [Ca^2+^] synchronization between a cell pair, as previously assumed^25^. Theoretically, it is common to observe unsynchronized node pairs^48^, though these networks tend to be relatively homogeneous, require symmetry, and rarely transmit information via a gradient^49^. Gap junctions facilitate ion passage between *β*-cells to synchronize electrical oscillations. Therefore, it makes sense that long-range synchronization requires a chain of highly conductive gap junctions (**Figure 4**), and that islet synchronization is decreased when gap junctions are removed (**Figure 5,6**). However, gap junctions are not the primary factor driving electrical activity. In *β*-cells, glucose metabolism drives electrical activity and [Ca^2+^] elevations. The cellular membrane conductance, driven by glucose metabolism, determines the electrochemical gradient along the gap junction, and thus the coupling current which drives synchronization^26,50^. Furthermore, gap junctions have a large role in repolarizing cells back to resting membrane potential^26^.

In answering question 3, we concluded that removal of edges in the structural network decreases the functional network efficiency and clustering. Decreased gap junction coupling^12,41–44^ and synchronization^12,41–44^ occur in type1 and type2 diabetes, but it is unclear whether altered synchronization is completely determined by altered gap junction coupling. Our computational and experimental results show that as structural edges were removed, the network efficiency and clustering decreased (**Figures 5,6**), implying that the islet loses efficient signal propagation and ability to synchronize. This in part explains decreased islet function in diabetes. However, neither clustering nor global efficiency could be explained by randomness (**Figure 6**). According to experiments^16^, only highly ordered networks will show zero overlap with randomness between the clustering and efficiency statistics. If gap junctions were solely responsible for variations in islet synchronization, a decrease in gap junction coupling would cause the functional network to appear more random. Instead, our findings indicate that *β*-cells are preferentially, not randomly, synchronized even in the presence of very low gap junction coupling. This is consistent with synchronization being driven by intrinsic cellular properties, such as metabolic activity. Thus while, removal of gap junctions is detrimental to proper islet function, synchronization is not directly indicative of structural connections. Further, perturbations in cellular metabolic activity, such as occurs in diabetes^51^ will have a significant influence on islet synchronization.

Our computational model is based on careful physiological recordings and has been previously validated^35–37^. Nevertheless, it will still be limited in describing *β*-cell dynamics. However, because our results are internally derived, where the functional and structural networks were obtained from the same model, our results are not dependent on the models’ ability to reflect islet dynamics. This fact also allows our methods to translate to any oscillatory network with similar characteristics. The choice of threshold (*R_th_*) is influential in creating the functional network and should be carefully considered. We used thresholds to match that of the prior experimental results^22,23,28^. The computational threshold was very high (*R_th_* = 0.9995) as the computational model did not include noise, so that synchronization was more acute. However, the trends in our results were consistent across thresholds (**Supp. 1**). In the computational model we also analyzed only fast electrical oscillations, with frequency <1 minute^52^, as in experimental measurements^23^. A similar analysis should consider slower metabolic oscillations and factoring in the (currently debated) mechanism by which metabolic oscillations may be synchronized across the islet. Finally, the islet also contains *α*-cells, *δ*-cells, and other cell types which can influence *β*-cell dynamics via paracrine mechanisms^53–55^. Our study only analyzes *β*-cells, as performed in other network studies which we compare^22,23,27^. However, we cannot exclude long-range structural projections from *δ*-cells^56,57^ and that *δ*-cells may be gap junction coupled^59^. Future analysis should include other cell types, which may provide valuable insight how their interactions can shape the overall islet response.

The islet is one of many biological systems where network theory has proved useful^60–62^. Therefore, the understanding that synchronization can give insight into cellular properties that drive excitability of the cell can be useful in many systems. Our results also show that forming structurally-based conclusions about a network using the functional network should be done with caution (we refer to^20^).

To summarize, we investigated the relationship between the synchronization-based functional network, the gap junction structural network, and *β*-cell metabolic activity in the islets of Langerhans. We concluded with three major points. *First*, highly connected subpopulations of *β*-cells, including *β*-cell hubs, are defined by glucose metabolism more than gap junctions. *Second*, metabolic similarities rather than gap junction connections drive edges in the functional network of the islet. *Third*, removal of edges in the structural network decreases global communication efficiency and local connectivity: decreasing gap junction conductance causes the functional network to become more sparse and less small world-like, which may have pathophysiologic implications. These results provide insight into the relationship between function and structure in biological networks and caution against using functional networks to make conclusions concerning the underlying structure and structural changes.

## Methods

### Calcium imaging, FRAP, and NAD(P)H

#### Animal care

Male and female mice were used under protocols approved by the University of Colorado Institutional Animal Care and Use Committee. β-cell-specific GCaMP6s expression (β-GCaMP6s) was achieved through crossing a MIP-CreER (The Jackson Laboratory) and a GCaMP6s line (The Jackson Laboratory)^63^. Genotype was verified through qPCR (Transetyx, Memphis, TN). Mice were held in a temperature-controlled environment with a 12 h light/dark cycle and given continuous access to food and water. CreER-mediated recombination was induced by 5 daily doses of tamoxifen (50mg/kg bw in corn oil) delivered IP.

#### Islet isolation and culture

Islets were isolated from mice under ketamine/xylazine anesthesia (80 and 16 mg/kg) by collagenase delivery into the pancreas via injection into the bile duct. The collagenase-inflated pancreas was surgically removed and digested. Islets were handpicked and planted into the glass-bottom dishes (MatTek) using CellTak cell tissue adhesive (Sigma-Aldrich). Islets were cultured in RPMI medium (Corning, Tewksbury, MA) containing 10% fetal bovine serum, 100 U/mL penicillin, and 100 mg/mL streptomycin. Islets were incubated at 37C, 5% CO2 for 24-72 h before imaging.

#### Imaging

An hour prior to imaging nutrition media from the isolated islets was replaced by an imaging solution (125 mM NaCl, 5.7 mM KCl, 2.5 mM CaCl2, 1.2 mM MgCl2, 10 mM HEPES, and 0.1% BSA, pH 7.4) containing 2 mM glucose. During imaging the glucose level was raised to 11mM. Islets were imaged using either a LSM780 system (Carl Zeiss, Oberkochen, Germany) with a 40× 1.2 NA objective or with an LSM800 system (Carl Zeiss) with 20× 0.8 NA PlanApochromat objective, or a 40× 1.2 NA objective, with samples held at 37C.

For [Ca^2+^] measurements GCaMP6s fluorescence was excited using a 488-nm laser. Images were acquired at 1 frame/s at 10-20 m depth from the bottom the islet. Glucose was elevated 3 minutes after the start of recording, unless stated otherwise.

NAD(P)H autofluorescence and [Ca^2+^] dynamics were performed in the same z-position within the islet. NADH(P)H autofluorescence was imaged under two-photon excitation using a tunable mode-locked Tisapphire laser (Chameleon; Coherent, Santa Clara, CA) set to 710 nm. Fluorescence emission was detected at 400–450 nm using the internal detector. Z-stacks of 6–7 images were acquired spanning a depth of 5 μm. First the NAD(P)H was recorded at 2 mM glucose, then the [Ca^2+^] dynamics was recorder at 2 mM and during transition to 11 mM glucose. After the [Ca^2+^] wave was established, the NAD(P)H was recorded at 11 mM glucose.

C×36 gap junction permeability and [Ca^2+^] dynamics were performed in the same z-position within the islet, with gap junction permeability measured using fluorescence recovery after photobleaching (FRAP), as previously described. After [Ca^2+^] imaging, islets were loaded with 12 mM Rhodamine-123 for 30 min at 37C in imaging solution. Islets were then washed and FRAP performed at 11mM glucose at room temperature. Room temperature was used because at this temperature the Rhodamine-123 travels between the cells only through the Cx36 gap junctions, versus at 37 C it can permeate a cell membrane. Rhodamine-123 was excited using a 488-nm laser line, and fluorescence emission was detected at 500–580 nm. Three baseline intensity images were initially recorded. A region of interest was then photobleached achieving, on average, a 50% decrease in fluorescence, and images were then acquired every 5-15 s for 15-30 min.

#### Analysis of [Ca2+] dynamics

Pearson-product-based network analysis presented in Figure 2 was performed as previously reported ^22^. [Ca^2+^] time courses were analyzed during the second-phase [Ca^2+^] response when the slow calcium wave was established. The Pearson product for each cell pair islet islet was calculated over each time point, and the time-average values were computed to construct a correlation matrix. An adjacency matrix was calculated by applying a threshold to the correlation matrix. The same threshold of 0.9 was applied to all islets. All cell pairs with a non-zero values in the adjacency matrix were considered to have a functional edge. The percent of edges was calculated with respect to the maximum number of edges per cell in each individual islet. For example, if a most connected cell possessed max=10 edges, and other cells had 1, 3, … 7 edges – then the % were: 10%, 30%, … 70%.

### Prior calcium imaging first presented in^1^

#### Islet isolation

Islets were isolated as described in Scharp et al.^64^ and Stefan et al.^65^ and maintained in Roswell Park Memorial. Institute medium containing 10% fetal bovine serum, 11 mM glucose at 37C under humidified 5% CO2 for 24–48 h before imaging.

#### Imaging islets

Isolated islets were stained with 4 mM Fluo-4 AM (Invitrogen, Carlsbad, CA) in imaging medium (125 mM NaCl, 5.7 mM KCl, 2.5 CaCl2, 1.2 mM MgCl2, 10 mM HEPES, 2mM glucose, 0.1% bovine serum albumin, pH 7.4) at room temperature for 1–3 h before imaging. Islets were imaged in a polydimethylsiloxane (PDMS) microfluidic device, the fabrication of which has been previously described in Rocheleau et al^66^, which holds the islet stable for imaging and allows rapid reagent change, such as varying glucose stimulation or adding gap junction inhibitors. Fluo-4 fluorescence is imaged 15 min after a step increase in glucose from low (2 mM) to high (11 mM). High speed imaging is performed on an LSM5Live with a 203 0.8 NA Fluar Objective (Zeiss, Jena, Germany) using a 488 nm diode laser for excitation and a 495 nm long-pass filter to detect fluorescence emission. The microfluidic device is held on the microscope stage in a humidified temperature-controlled chamber, maintained at 37 C. Images were acquired at a rate of 4–6 frames per second, with average powers at the sample being minimized to, 200 mW/cm2.

#### Analysis of Ca2+ Imaging data

We present data from 11 Islets from 6 WT mice, 11 Islets from 11 heterozygous C×36 +/− knockout mice, and 14 Islets from 3 homozygous C×36 -/- knockout mice. We extracted cell calcium dynamics by visually identifying and circling all cells islet. We assumed that the pixels within a cell should be well coordinated, so we removed any pixels whose dynamics were not within 5-10 STD of the average. This usually resulted in the removal of 1-5 pixels on the edge of the cell boundary.

#### Threshold

Previous studies have shown that wild type Islets should have a degree distribution that is roughly linear when plotted on a log-log plot and average degree of either 8 or between 5-15^22,23,28,67^. To determine R_th_ that best satisfied both of these findings, we utilized constrained optimization matlab algorithm fminsearchbnd^68^ to find the R_t_h_opt_ which maximized the goodness of fit to a power law distribution, while forcing 5 ≤ k_avg_ ≤ 15 for each WT islet constrained optimization. The average optimal threshold (*R*_th_ = 0.90).

#### Average distance between connected cells

The average distance between connected cells calculated the total number of pixels between the center of two connected cells. The average distance was expressed as the normalized distance between connected cells and the average distance between all cells in the islet to control for image and Islet size.

### Computational Model

This ordinary differential equation model has been described previously ^26^ and validated on experimental studies ^34–37^. This model has been shown to accurately describe experimental findings concerning spatial-temporal dynamics ^37^, the relationship between electrical coupling and metabolic heterogeneity ^36^, the relationship between electrical heterogeneity and electrical activity ^35^, and the influence of excitability parameters such as rate of glucose metabolism (k_glyc_) and K_ATP_ channel opening kinetics on diabetes mutations ^34,36^. It is based on a single *β*-cell electrophysiologic model ^69^, in which the membrane potential (*V_i_*) for *β-cell_i_* is solved for each time step using eq 1a. We created a 1000 *β*-cell network and electrically coupled any cell pairs within a small distance from each other. We chose 1000 *β*-cell because most species (including human and mouse) contain on average 1000 *β*-cells in an islet ^70^ and it is the number which has been validated experimentally. The coupling current, *I_coup_* is determined by the difference in voltage between the cell pair and the average coupling conductance (*g_coup_*) between them (1b). All code was written in C++ and run on the SUMMIT supercomputer (University of Colorado Boulder). All simulations are run at 11mMol glucose. The membrane potential (*V_i_*) for *β-cell_i_* is solved for using the ODE:

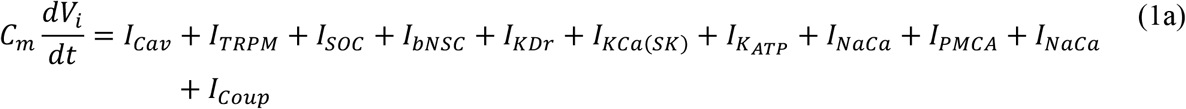

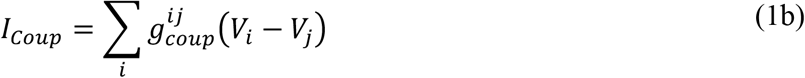

The average number of gap junction connections per cell was 5.24. Cellular heterogeneity was introduced by assigning randomized metabolic and electrical parameter values to each cell based on their distributions previously determined by experimental studies. For example, coupling conductance (g_coup_) was assigned so that the Islet average is *g_coup_* = 0.12*nS* and glucokinase (*k_glyc_*), the rate-limiting step in glycolysis, was assigned so that the average *k_glyc_* = 1.26 * 10^−04^ *sec*^−1^. We ran the simulation for 500 seconds and only the 2^nd^ phase was analyzed, in concordance with ^23^. All the results presented are based on five different Islets with randomized parameter values. *β*-cell heterogeneity was simulated by randomizing parameters including coupling conductance (*g_coup_*) and metabolic rate (*k_glyc_*) according to experimental results. To explore the effects of coupling conductance on the functional network, we altered Islet average coupling conductivity to *g_coup_* = 0.06 *nS* and *g_coup_* = 0.03 *nS*, and then assigning each cell a randomized *g_coup_* based on the new target average. The gap junction conductance (g_coup_) for any cell was calculated as the sum of the average conductivity (g_coup_) between the cell of interest and any cells it shares a gap junction connection with:

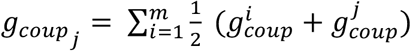

### Network Analysis

#### Creating Functional Network from simulated Islet

The methodology was based on that previously defined in ^22^. First, the correlation coefficient between each cell was calculated using corr() function in matlab, which follows equation: 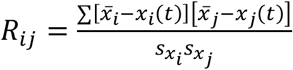. Next, a threshold (*R_th_*) was defined to determine whether each cell pair is ‘synchronized’ or ‘not synchronized’. For computational experiments, the threshold was chosen such that the network roughly followed a power law distribution, as predicted in ^22,23^. Unless otherwise noted, *R_th_* = 0.9995 for computational analysis.

#### Creating the Metabolic network

Because gap junctions enforce localization onto the analysis, we looked at cell pairs whose Pythagorean distance was ≤ 15% within the islet. Within this sample space, we looked at cell pairs whose average kglyc was higher than the Islet average (**Figure 4b**, green) (0.5 * |*k_glyc_i__* + *k_glyc_j__* | > *average*(0.5 * *k_glyc_i__* + *k_glyc_j__*, | ∀_*ij*_)), and cell pairs whose k_glyc_ was more similar than the islet average (*|*k_glyc_i__*, – *k_glyc_j** | < *average*(|*k_glyc_i__*, – *k_glyc_j__* | ∀_*ij*_)). We chose these parameters such that the *Pr*(*Sync|GJ*) = *Pr* (*Sync|Met*), which acts as a control allowing us to directly compare *Pr*(*GJ|Sync*) to *Pr*(*Met|Sync*).

#### Average degree

The degree *k_i_* was calculated by counting the number of connections for *cell_i_* and averaging k over all cells islet, then normalized to the Islet size to remove any size dependence (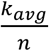, *where* ≡ *islet size*).

#### Degree Distribution

The degree distribution was calculated by first calculating the degree of each cell by taking the column sum of the adjacency matrix. Then each cell degree was normalized by dividing by the maximum degree of the islet. The histogram was then calculated using GraphPad Prism.

##### Hub identification

*In accordance with Johnston et al., any cell with more than 60% islet, calculated by the degree distribution, was considered a hub cell*.

#### Probabilities

We first created the functional network and structural (GJ related probabilities) or metabolic network (metabolism). We then calculated the probability that two cells were synchronized by:

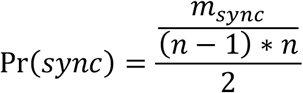

Where *m_sync_* = number of edges in the synchronized network, and *n* = number of nodes. Similarly, we calculated the probability that two cells were either gap junction coupled or metabolically related using

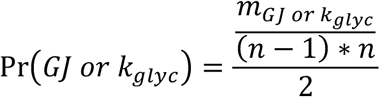

Where *m_GJ or k_glyc__* is the number of edges in the gap junction or metabolic network.

To find the probability that a cell pair was both synchronized and GJ or metabolically connected, we calculated

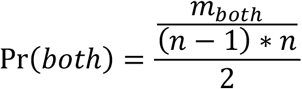

Where *m_both_* is any edge that exists in both matrices.

Finally, we calculated conditional probabilities by:

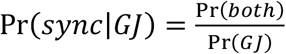

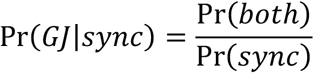

These quantities were calculated separately for each Islet seed and then averaged.

### Network topology analysis

#### Shortest weighted path length

The gap junction network was weighted using the inverse of g_coup_ or k_glyc_ between *Cell_i_* and 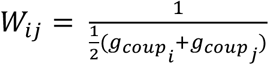 or 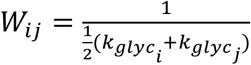. This is done because the shortest path length algorithm views weights as “resistance” and finds the path of least resistance. The shortest weighted path between every cell was calculated using Johnson’s algorithm ^71^. Cell pairs were then categorized as synchronized or not synchronized if their correlation coefficient was < *R_th_*. The average for synchronized and non-synchronized cells was calculated over each islet for each distance. To normalize over distance, each data point was divided by the average of the non-synchronized and synchronized islets for the given distance.

#### Clustering Coefficient

The clustering coefficient represents the ‘cliquishness’ of the network. This is defined by ^22^ as the “number of existing connections between all neighbors of a node, divided by the number of all possible connections between them.” This was calculated by making a subgraph of each cell’s connections and counting the number of connections between those cells. For example, if A is connected to B, C, and D, and B and C are connected but D is not connected to any other cell (see matrix). Then the clustering for cell A is 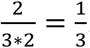. Each node is a assigned a clustering coefficient *C* such that:

| Subgraph<br>- A | B | C | D |
| --- | --- | --- | --- |
| B |  | 1 | 0 |
| C | 1 |  | 0 |
| D | 0 | 0 |  |

The average clustering coefficient is 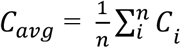.

#### Average Shortest Path Length

Shortest path length was calculated with matlab function graphallshortestpaths(). This function uses the Johnson’s algorithm ^71^ to find the shortest path between every pair of cell islet. For example, the path length between *cell_i_*. and *cell_j_* is 1 if they are directly synchronized, or 2 if *cell_i_*. is not synchronized with *cell_j_* but each is synchronized with *cell_k_*. To compensate for highly sparse network, any non-connected node was given a characteristic path length of *n* + 1. Finally, the characteristic path length (L), or average path length, was expressed as the sum of all path lengths normalized to total possible connections (size*(size-1)).

The normalized average shortest path length (Supplemental 2d) is therefore calculated as 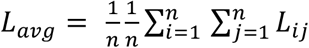. To compensate for any non-connected cell, whose path length *L_ix_* = ∞, where *x* is any cell islet, we set the path length for non-connected cells to *L_ix_* → *n* + 1 where *n* is the number of cells islet.

#### Global Efficiency

The global efficiency is related to the inverse of global path length ^40^.

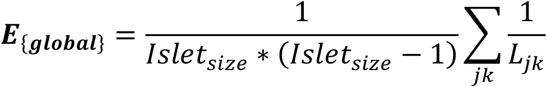

length using 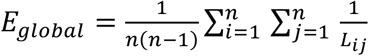. Because *E_global_* =0 for a non-connected cell, the disconnected network is naturally compensated for.

#### Random Networks

Random networks were created using an Erodos Renyi approach ^72^. For each islet, we created 1000 random networks with equal number of nodes, edges, and average degree as the islet of interest. The probability that two cells were connected was given by 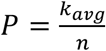. For each cell pair within the islet, a random number generator created a number from 0-1. If that number was below P, whose cells were connected by an edge.

#### Small World properties

For each islet, small world parameters *S_E_* and *S_L_* were calculated 1000 times based on the 1000 random networks.

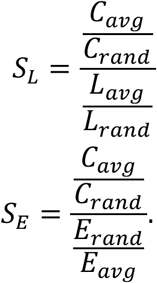

#### Local Efficiency

Local efficiency is calculated by first extracting a subgraph based on the connections for an individual cell. Within that cell, the efficiency is calculated by:

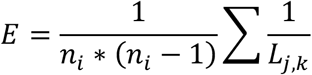

Where *n_i_* is the number of nodes that an individual node is connected to. This is done for every node in the network and all efficiencies are averaged to present the local efficiency of the network.

### Significance Testing

Significance tests are done in Prism (GraphPad).

The computational model is run on 5 seeds, therefore all significance tests are paired.

Figure 1e-g, Figure 4e&h, and Figure 5c-f are paired, two tailed t-tests.

Figure 3e,f are repeated measures paired one-way ANOVA with multiple comparison using Tukey’s multiple comparison.

Figure 4i,j are multiple paired t-tests with Bonferroni-Dunn adjustment.

All significance tests in figure 6 are unpaired ordinary one-way ANOVA tests.

Figure 7 c-j are ordinary paired one-way ANOVA with Tukey multiple comparison test.

Supplemental 1b and 2b, Figure 4 i and j and Figure 5i: To compare, we used a paired T-test with Bonferroni multiple comparisons corrections. Since we there were 15 cell distances, we set the significance threshold *α* = 0.003. For convenience, we present asterisks next to significant P-values. Supplemental 4b and c are two way ANOVA with Sidak’s multiple comparisons.

## Acknowledgements

Richard KP Benninger (University of Colorado) is the guarantor of this work and, as such, had full access to all the data in the study and takes responsibility for the integrity of the data and the accuracy of the data analysis. We thank David W Piston (Washington University St Louis) with whom data presented in figure 6 was previously acquired. We thank Aaron Clauset for helpful conversations and advice. All authors acknowledge that no conflict of interest exists.

## Author Contributions

Conceptualization: JKB, RKPB, VK

Methodology: JKB, RKPB, VK, JMD

Investigation: JKB, VK

Visualization: JKB

Supervision: RKPB

Writing—original draft: JKB

Writing—review & editing: JKB, RKPB, VK, DJA

## Data and materials availability

Raw microscopy imaging data to the EMBL-EBI-supported BioImage Archive. Analysis code, Model code, and Simulated Data will be available via GitHub at https://github.com/jenniferkbriggs/Functional_and_Structural_Networks.git

