## Supplementary material for "Beta-cell Metabolic Activity Rather than Gap Junction Structure Dictates Subpopulations in the Islet Functional Network": Supp.

**This PDF file includes:**

Figs. S1 to S7

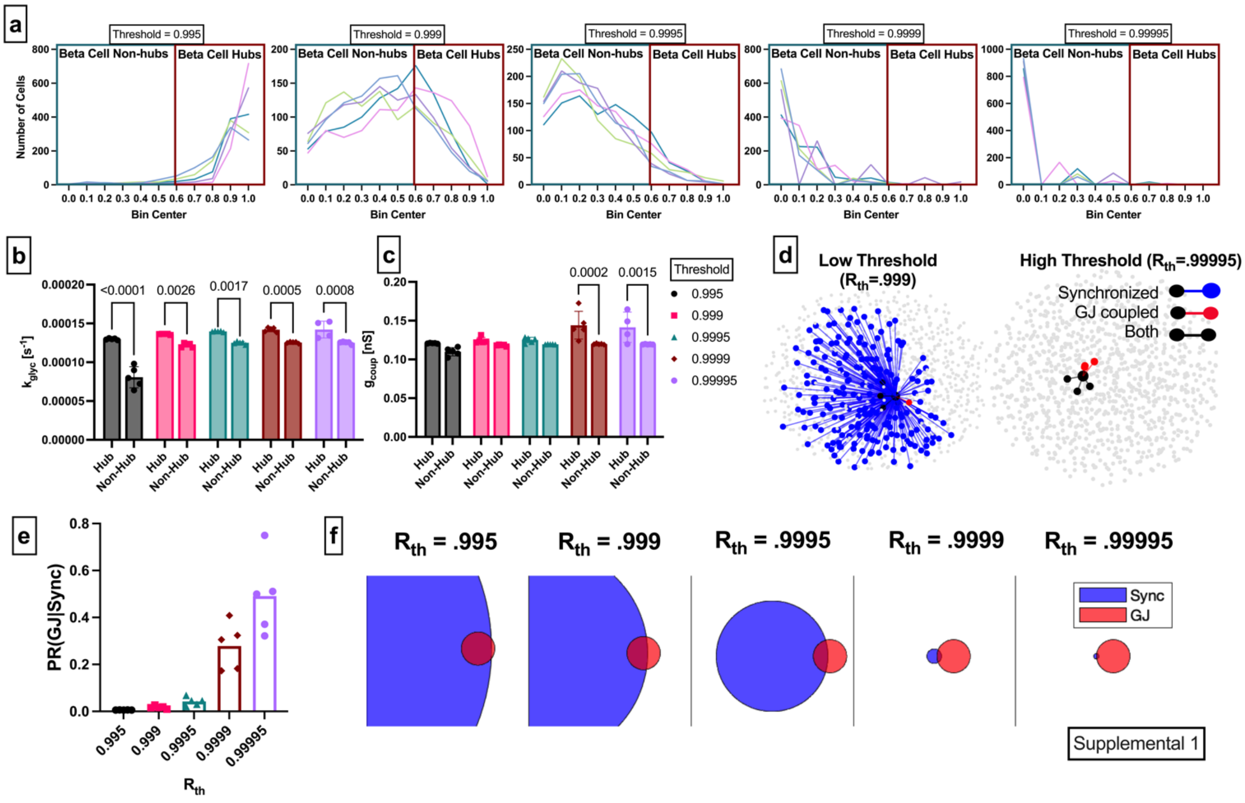


**Fig. S1. Network sensitivity to threshold**

**a:** Degree distribution of five simulated islets with five different thresholds. **b:** $k_{glyc}$ comparing hubs and non hubs across five different thresholds. **c:** $g_{coup}$ comparing hubs and non hubs across five different thresholds. Significance is assessed using Tukeys Multiple Comparisons. **d:** Islet with representative cell connections calculated by a lower threshold ($R_{th} = 0.999$) (**right**) and higher threshold ($R_{th} = 0.99995$) **(left)**. Synchronized cell pairs are shown in blue, gap junction pairs are shown in red and both are shown in black. **e:** Probability of a gap junction connection given synchronization for the five thresholds. **f**: Venn diagrams for the probability of synchronization (blue) and gap junction connection (red) with the overlapping area representing cell pairs that were both synchronized and gap junction coupled.

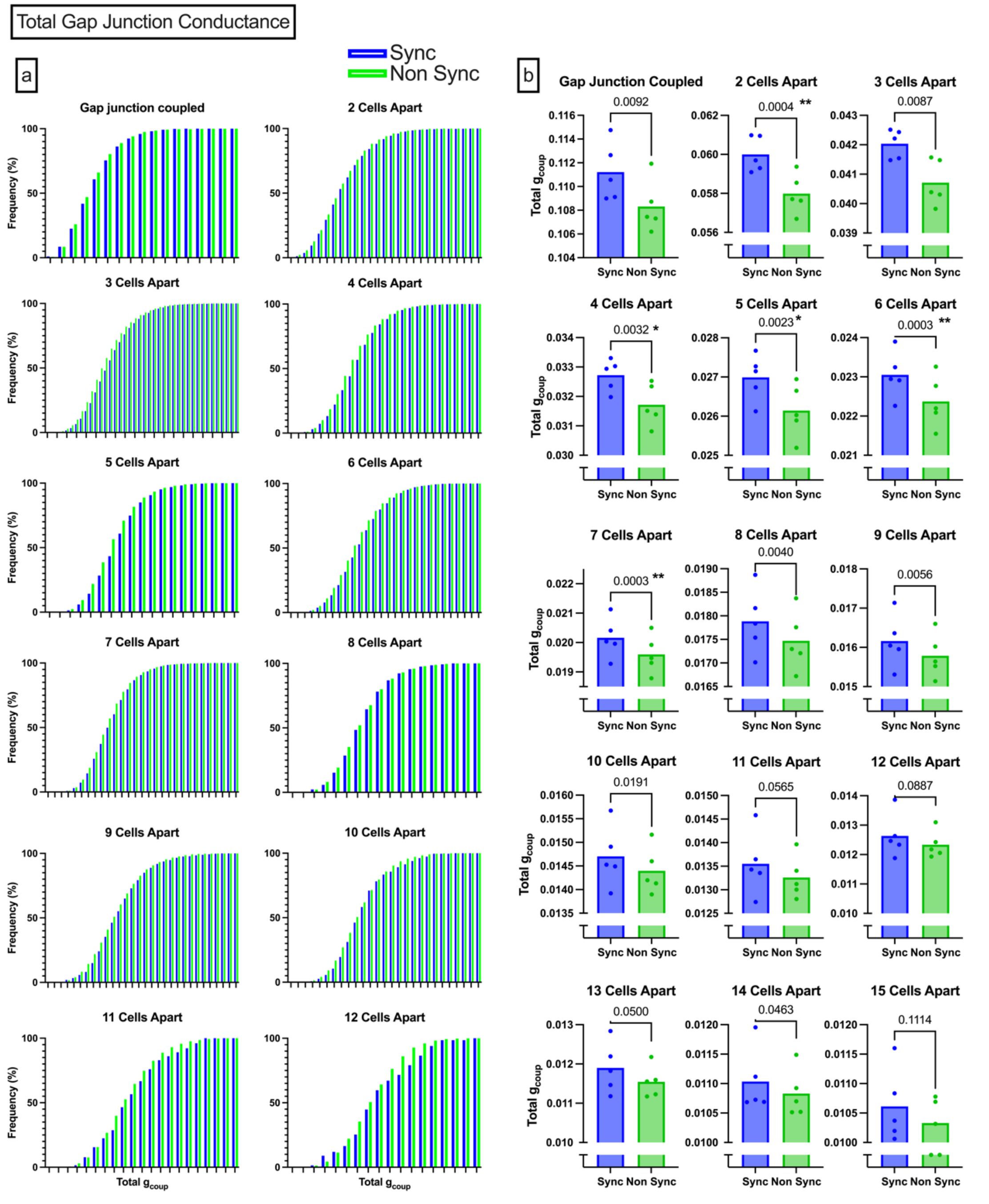


Supplemental 2

**Fig. S2: Paths that maximized gap junction conductivity.**

**a**: Cumulative distribution functions of total conductance between two cells from 1-12 cells apart. Synchronized CDF is shown in blue, non-synchronized CDF is shown in green. **b**: Bar charts comparing the average conductance for synchronized and non-synchronized cells. P values are not corrected, but significance (denoted by asterisks) is determined using a Bonferroni-corrected paired T-test. Since we there were 15 cell distances, significance threshold was set at $\alpha = 0.003$.

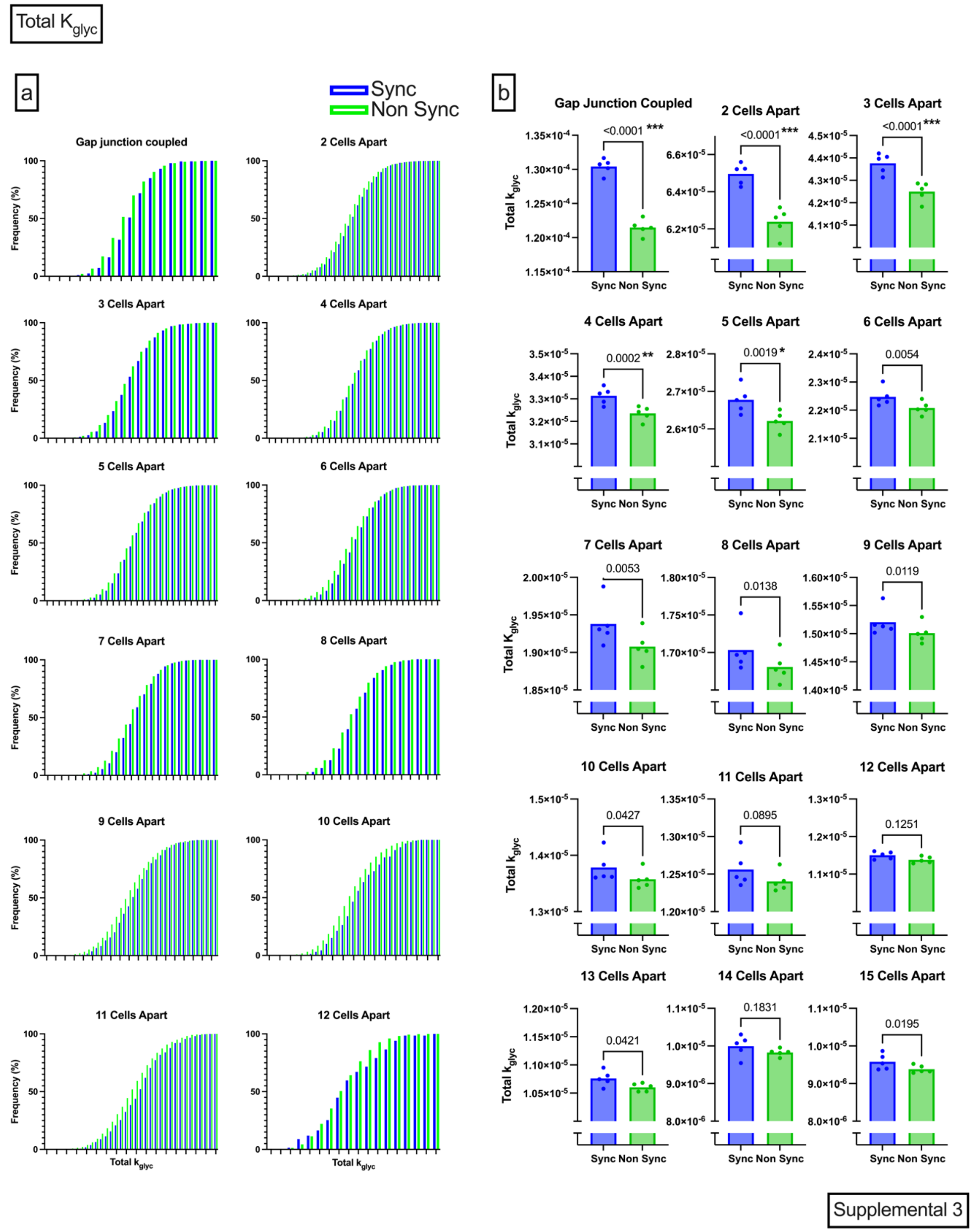


**Fig. S3. Paths that maximized metabolic rate.**

**a:** Cumulative distribution functions of total $k_{glyc}$ between two cells from 1-12 cells apart. Synchronized CDF is shown in blue, non-synchronized CDF is shown in green. **b:** Bar charts comparing the average conductance for synchronized and non-synchronized cells. P values are not corrected, but significance (denoted by asterisks) is determined using a Bonferroni-corrected paired T-test. Since we there were 15 cell distances, significance threshold was set at $\alpha = 0.003$.

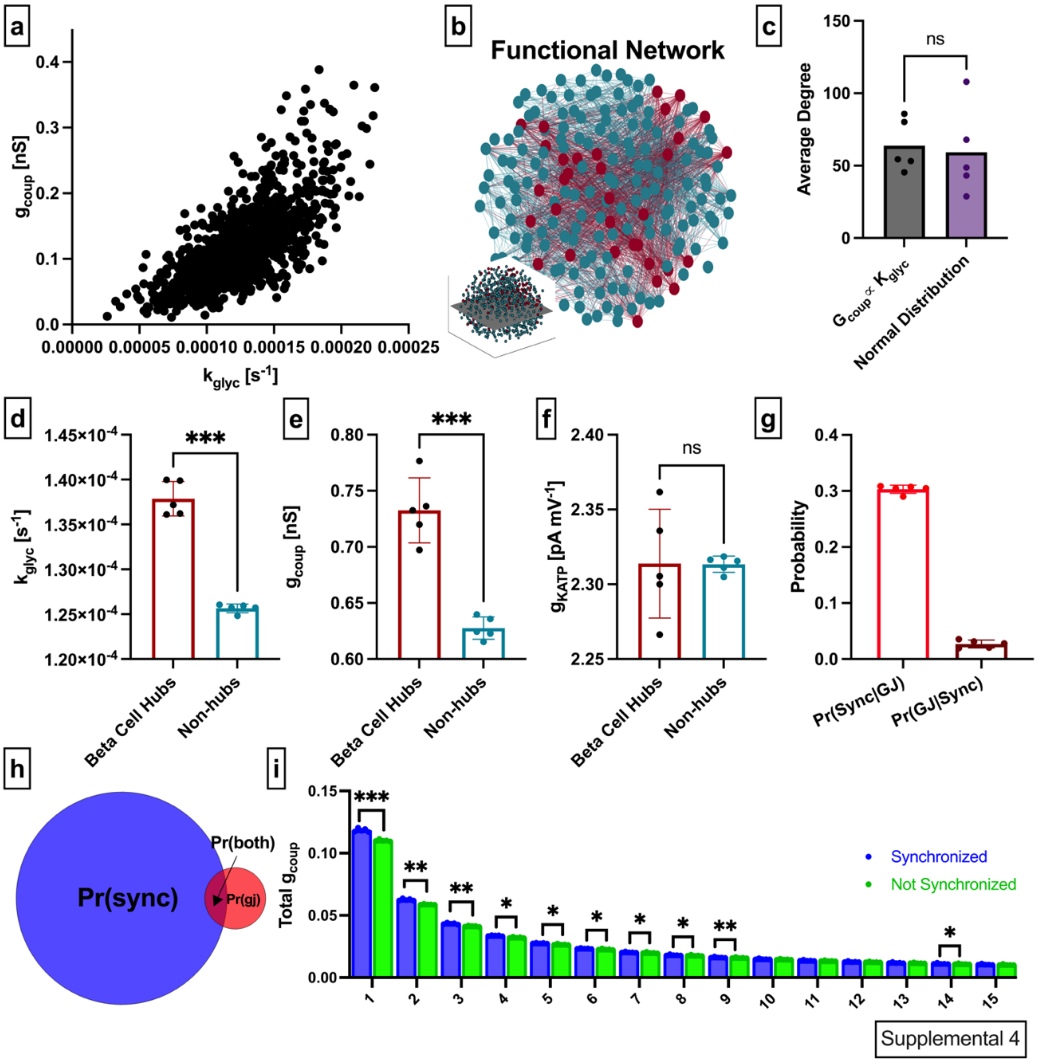


**Fig. S4: Assessment of functional and structural network overlap in a correlated distribution of glucose metabolism and gap junction conductance**

**a:** Distribution where $g_{coup}$ is correlated with $k_{glyc}$, such that cells with high metabolism also have high gap junction coupling. **b:** 2-dimensional slice of the islet, with lines (edges) representing functional connections between synchronized cells. $\beta$-cell hubs are indicated in red. Slice is taken from middle of the islet (see inset). **c:** Average degree comparing proportional distribution shown in a and normal distribution (Figure 1), where there is no relationship between $k_{glyc}$ and $g_{coup}$ **d:** Average rate of glucose metabolism ($k_{glyc}$) parameter values compared between hub and non-hub, retrospectively analyzed. **e:** as in d for gap junction conductance ($g_{coup}$). **f:** as in d for maximum conductance of ATP sensitive potassium channel ($g_{KATP}$). **g:** probability of synchronization given a cell pair is gap junction coupled P(Sync|GJ), and probability that a cell pair which is gap junction coupled, is synchronized P(GJ|Sync). **h:** Venn diagram showing overlap between synchronized cell pairs and gap junction connected cell pairs. **i:** Total $g_{coup}$ for synchronized and not synchronized cell pairs organized by cell distance. Shaded area in h is proportional to indicated probability. Significance in c-g was determined by 2-tailed paired Students t-test. Significance in i was assessed by paired t-tests with Bonferroni correction. $*P \leq 0.05$, $**P \leq 0.01$, $***P \leq 0.001$.

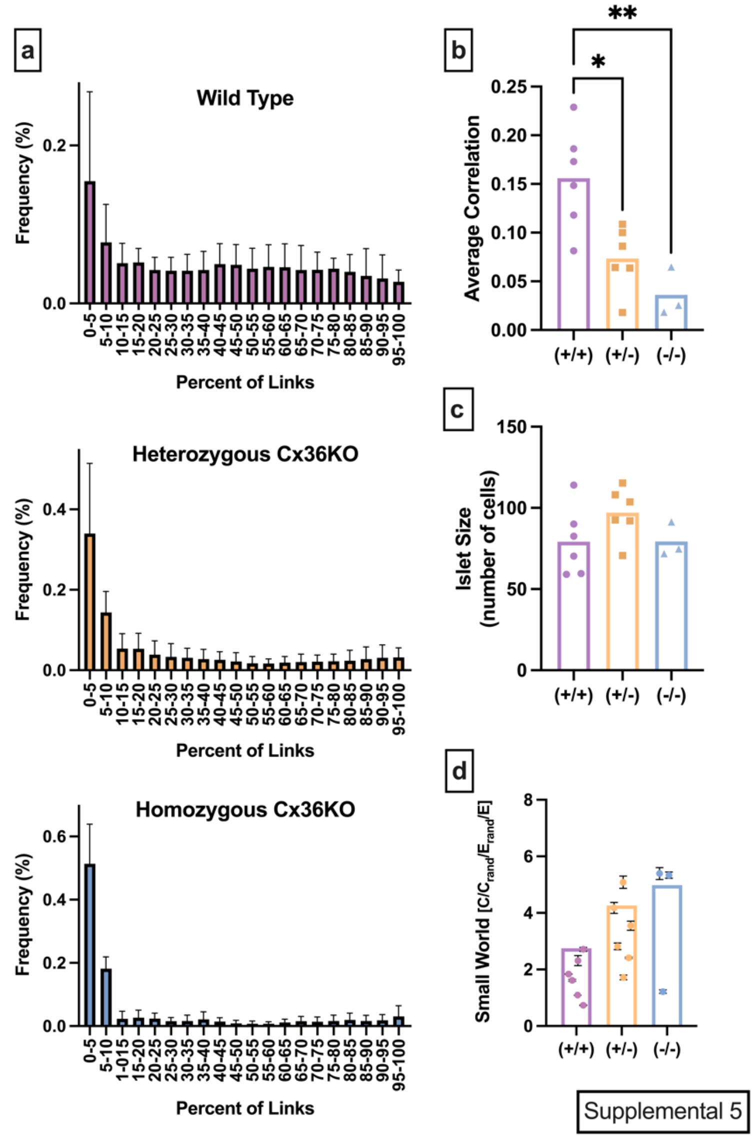


**Fig. S5: Alternative Network Metrics for Calcium Islets from Cx36-KO mice**

**a**: Averaged degree distribution histograms for all Islets analyzed. Error bars indicate standard deviation. Purple (**top**) histogram shows the distribution of connections for wild type ($Cx36^{+/+}$) islets. Orange (**middle**) shows the distribution of connections for heterozygous Cx36 knockout ($Cx36^{+/-}$) islets. Blue (**right**) shows the distribution of connections for homozygous Cx36 knockout ($Cx36^{-/-}$) islets. **b**: Average Pearson cross-correlation coefficient, between all islet in the islet. **c:** Number of cells in each islet. **d**: Small world-ness of a network calculated using global efficiency. 1000 random networks per islet were created to determine uncertainty of each data point (shown in black). Significance, or lack thereof is determined by an ordinary one-way ANOVA and Tukey Multiple comparison test. $*P \leq 0.05$, $**P \leq 0.01$.

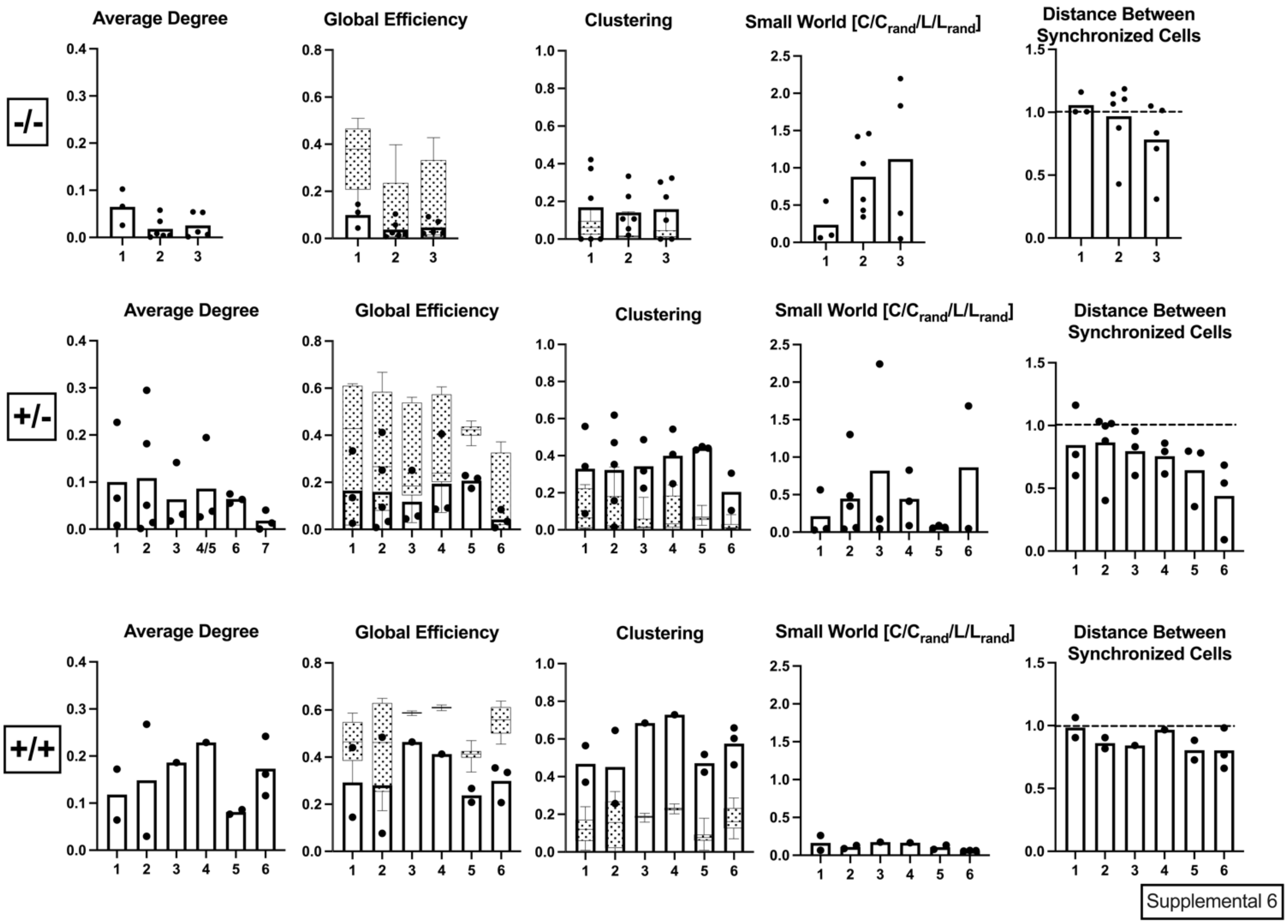


**Fig. S6: Network Metrics assessed by Islets**

This figure shows similar information as that of Figure 6, but the data points are individual islets, and the bars are those Islets grouped by mouse. Genotypes from top to bottom: homozygous Cx36 knockout ($Cx36^{-/-}$) islets, heterozygous Cx36 knockout ($Cx36^{+/-}$) islets, wild type ($Cx36^{+/+}$) islets. Random network distributions are overlayed as box-and-whisker plots.

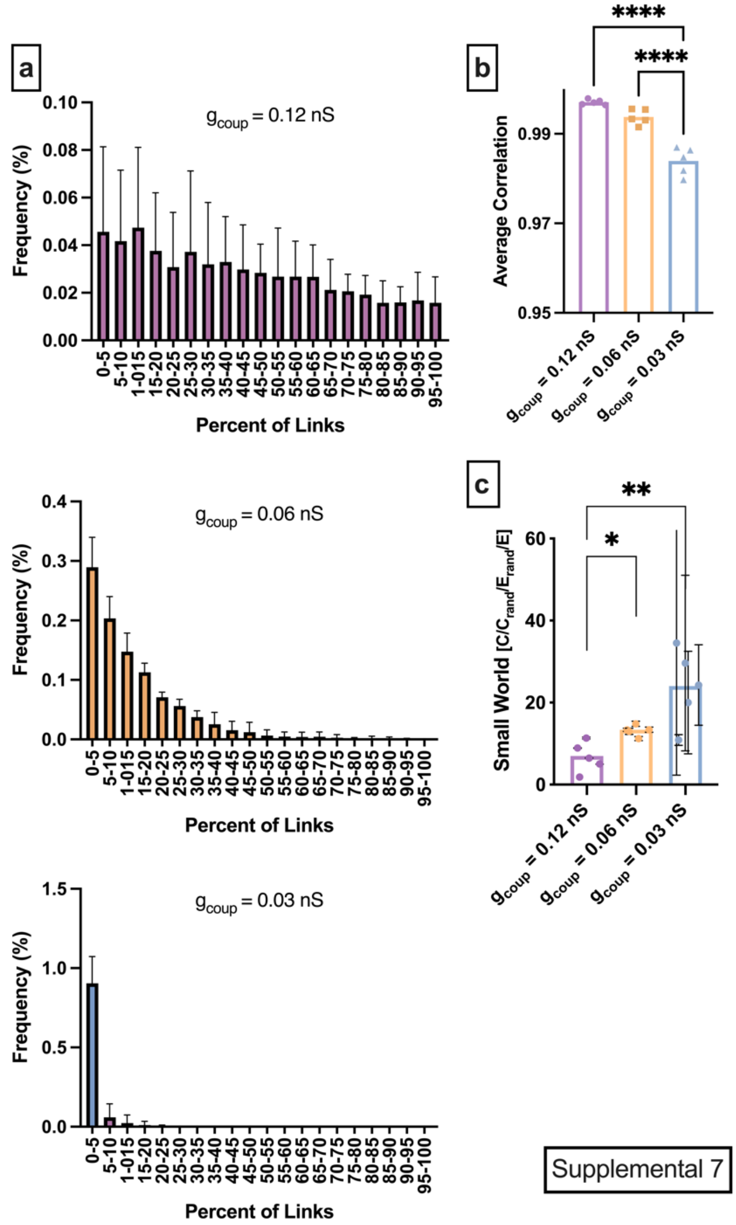


**Fig. S7: Alternative Network Metrics for Simulated Islet with Gap Junction Coupling Change**
**a**: Averaged degree distribution histograms for all Islets analyzed. Error bars indicate standard deviation. Purple (**top**) histogram shows the distribution of connections for fully coupled (0.12 nS) islets. Orange (**middle**) shows the distribution of connections for half coupled (0.06 nS) islets. Blue (**right**) shows the distribution of connections for homozygous quarter coupled (0.03 nS) islets. **b**: Average Pearson cross-correlation coefficient between all islets in the islet. **c:** Small world-ness of a network calculated using global efficiency. 1000 random networks per islet were created to determine the uncertainty of each data point (shown in black). Significance or lack thereof is determined by an ordinary one-way ANOVA and Tukey Multiple comparison tests. $*P \leq 0.05$, $**P \leq 0.01$
